# The organization of intracortical connections by layer and cell class in the mouse brain

**DOI:** 10.1101/292961

**Authors:** Julie A. Harris, Stefan Mihalas, Karla E. Hirokawa, Jennifer D. Whitesell, Joseph E. Knox, Amy Bernard, Phillip Bohn, Shiella Caldejon, Linzy Casal, Andrew Cho, David Feng, Nathalie Gaudreault, Charles R. Gerfen, Nile Graddis, Peter A. Groblewski, Alex Henry, Anh Ho, Robert Howard, Leonard Kuan, Jerome Lecoq, Jennifer Luviano, Stephen McConoghy, Marty T. Mortrud, Maitham Naeemi, Lydia Ng, Seung W. Oh, Benjamin Ouellette, Staci A. Sorensen, Wayne Wakeman, Quanxin Wang, Ali Williford, John W. Phillips, Allan Jones, Christof Koch, Hongkui Zeng

## Abstract

The mammalian cortex is a laminar structure composed of many cell types densely interconnected in complex ways. Recent systematic efforts to map the mouse mesoscale connectome provide comprehensive projection data on interareal connections, but not at the level of specific cell classes or layers within cortical areas. We present here a significant expansion of the Allen Mouse Brain Connectivity Atlas, with ∼1,000 new axonal projection mapping experiments across nearly all isocortical areas in 49 Cre driver lines. Using 13 lines selective for cortical layer-specific projection neuron classes, we identify the differential contribution of each layer/class to the overall intracortical connectivity patterns. We find layer 5 (L5) projection neurons account for essentially all intracortical outputs. L2/3, L4, and L6 neurons contact a subset of the L5 cortical targets. We also describe the most common axon lamination patterns in cortical targets. Most patterns are consistent with previous anatomical rules used to determine hierarchical position between cortical areas (feedforward, feedback), with notable exceptions. While diverse target lamination patterns arise from every source layer/class, L2/3 and L4 neurons are primarily associated with feedforward type projection patterns and L6 with feedback. L5 has both feedforward and feedback projection patterns. Finally, network analyses revealed a modular organization of the intracortical connectome. By labeling interareal and intermodule connections as feedforward or feedback, we present an integrated view of the intracortical connectome as a hierarchical network.

---

Cognitive processes and voluntary control of behavior originates in the isocortex. Understanding how incoming sensory information is processed, integrated with past experiences and current states to generate memories, percepts, and motor outputs requires knowledge of the anatomical patterns and rules of connectivity between cortical areas. Connectomes, complete descriptions of the wiring in a brain^1^, exist at different levels of spatial granularity (micro-, meso-, and macro-scale), each revealing different principles of brain organization. At the mesoscale^2^, connectivity is described at the level of cell populations, classes, or types. Recently, several mesoscale cortical connectomes have been produced, either through new systematic data generation^3–5^, or via expert collation of historical tract tracing literature^6–8^. Common organizational features of cortical connectivity in both rodent and macaque have been distilled from these datasets, often using graph theoretical approaches to describe network architecture^9,10^. For example, areas have unique patterns of connections (*i.e.,* a “fingerprint”), connection strengths follow a log-normal distribution spanning> 4 orders of magnitude^4,5^, and networks display high clustering coefficients, with some highly connected nodes (“hubs”)^4^. Finally, the organization of cortical areas is modular, with distinct modules corresponding to specific functions^3,11^.

Organizational schemes other than modular networks have previously been applied to cortical connections to explain information flow and processing. Specifically, the concept of a cortical hierarchy^12,13^ has been useful for understanding computational and architectural properties of the cortex and has inspired the development of neuronal network methods in machine vision^14^. These schemes (modular and hierarchical) are not mutually exclusive, and the actual organization of the cortex involves both types.

So far, the available mesoscale datasets are not clearly differentiated from macroscale connectivity in that the experiments and analyses focus on connections between areas, treating them as more or less homogenous regions. Data are often generated using tracers that cover entire cortical columns. In the first phase of the Allen Mouse Brain Connectivity Atlas, injections were intentionally placed at multiple depths within a cortical area in one experiment to infect all neurons across the cortical layers^4^. However, each cortical region is composed of a heterogeneous mix of distinct cell types. Perhaps the most characteristic feature of the isocortex is its organization into six layers. Within these layers, distinct cell types exist that can be further differentiated based on morphological, electrophysiological, and transcriptional properties^15–17^. Specific long-distance connectivity patterns are associated with excitatory cell populations or genetically-identified types in each layer^16,18^. Long-distance axon projections are commonly used to classify excitatory neurons into three main classes; intratelencephalic (IT), pyramidal tract (PT), and corticothalamic (CT). Axons from IT neurons project to both ipsilateral and contralateral cortex and striatum. PT neurons target subcortical structures, including those in the spinal cord, medulla, pons, and midbrain, and can send branches to ipsilateral cortex, striatum, and thalamus. CT neurons project to ipsilateral thalamus. IT neurons are found across all layers containing excitatory cells (L2/3, L4, L5, L6), while PT neurons locate to L5 and CT neurons to L6^19,20^. Projections from these major classes to different target regions suggests that they play distinct roles in information processing.

Experimental access to cortical cells in different layers and classes is feasible due to the generation of diverse Cre driver transgenic mouse lines^21–24^. By taking advantage of them, we significantly expanded upon our previously published Allen Mouse Brain Connectivity Atlas (http://connectivity.brain-map.org^4,25^) adding ∼ 1,000 new experiments in the cortex. Here we present this enhanced online resource of projection data from cortical cell classes defined by laminar position and brain-wide projection patterns. We first describe the macroscale organization of cortical connectivity into six network modules and the unique patterns of intracortical connectivity from each source. Then, we show the contribution of each layer-specific projection neuron class to the complete intracortical projection pattern for a given region. We observe diverse axon lamination patterns in cortical targets related to the layer of origin and projection neuron class labeled by each Cre line. These patterns are both similar to and different from previous anatomical patterns derived from anterograde tracing to define hierarchical position (feedforward and feedback) in the visual cortex of primates^12^ and rodents^26^. Using the cell layer/class-based projection patterns, we ordered 37 of 43 isocortical areas into hierarchical positions. We also ordered the network modules identified here; from bottom to top: somatomotor, temporal, visual, medial, anterolateral, and prefrontal cortex.

## Results

### Data generation and characterization of Cre driver lines for layer- and cell class-selective mapping

Our pre-existing pipeline for generation and quantification of projection mapping data across the entire brain^4^ was used in this study, with some modifications. In brief, in Oh et al., 2014 we used only wild-type C57BL/6J mice injected with a Cre-independent tracer, rAAV2/1.hSyn.EGFP.WPRE. Here, we used transgenic Cre driver mice. Most Cre mice were injected with a Cre-dependent rAAV tracer, rAAV2/1.pCAG.FLEX.EGFP.WPRE. A subset had a duplicate injection of an rAAV with a synaptophysin-EGFP fusion transgene in place of the cytoplasmic EGFP, rAAV2/1.pCAG.FLEX.SypEGFP.WPRE. Results from these two tracers were highly and significantly correlated for injections in close spatial proximity (**Supplemental Figure 1**). Thus, we included the SypEGFP datasets when indicated for analyses of connectivity patterns from given source areas. Following tracer injections, brains were imaged using serial 2-photon tomography^27^ (STP) at high x-y resolution (0.35 μm x 0.35 μm) every 100 μm, after which the images underwent QC and manual annotation of injection sites, followed by signal detection and registration to the Allen Mouse Brain Common Coordinate Framework (CCF), our fully annotated 3D volumetric reference atlas, for subsequent data analyses, data visualization, and public release to our web portal^28^.

Our goal for expanding on the Allen Mouse Brain Connectivity Atlas (http://connectivity.brain-map.org) was to create a comprehensive map of all inter-areal projections originating from neurons of different cell classes within a given source. Forty-nine Cre driver lines (**Supplemental Table 1**) entered the pipeline for cortical projection mapping after initial anatomical characterization of transgene expression across the brain using *in situ* hybridization^23^ (http://connectivity.brain-map.org/transgenic). These driver lines have either pan-layer Cre expression (e.g. Emx1-IRES-Cre), layer-selective Cre expression, or Cre expression driven by inhibitory neuron-specific promotors. Layer-selectivity data for 15 lines are summarized in the second row (“Layer”) of Figure 1a. Data used to determine layer-selectivity can be found in **Supplemental Figure 2**. Many Cre lines showed relatively even distribution of expression across the entire isocortex, but we also saw gradients and area-restricted expression patterns which were used to choose appropriate locations for tracer injections.

**Figure 1.**
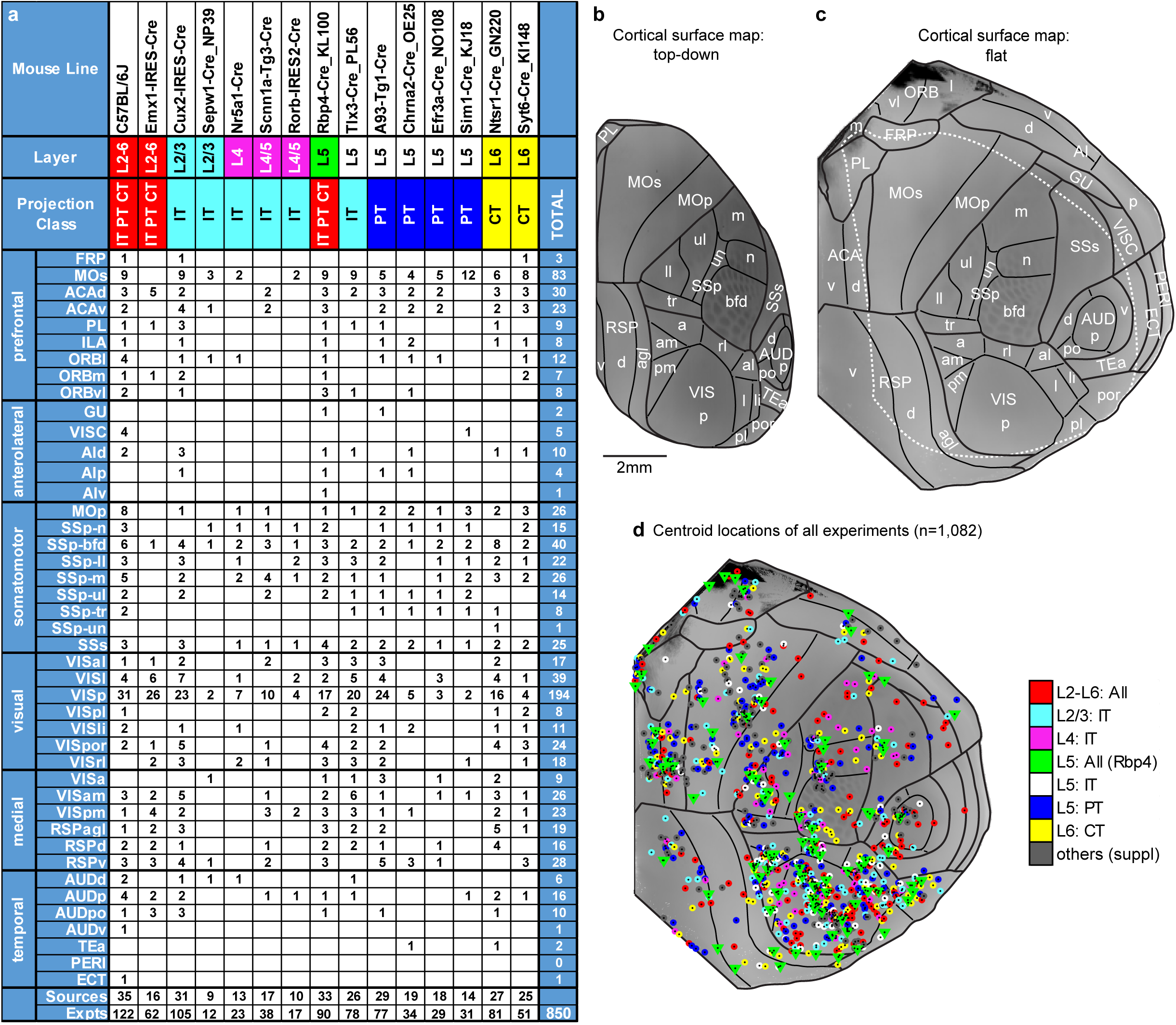
Systematic generation of cortical projection mapping data by area and mouse line. **(a)** Location and number of 850 tracer injection experiments across cortical areas and selected mouse lines. These 15 lines (C57BL/6J through Syt6-Cre_KI148) were used most extensively across cortical regions to map projections using Cre driver lines with expression preferentially in the layers and projection neuron classes indicated **(b)** Mouse isocortex is parcellated into 43 areas in the Allen CCF. The positions of most areas are visible in our standard top-down view of the right hemisphere cortical surface. This view is obtained by projecting the maximum density voxels from the average template brain, used to construct the CCF, along a curved coordinate system meant to match the columnar structure of the cortex (as opposed to a direct z-projection). **(c)** Areas that occupy very lateral, frontal and midline positions are better viewed in a flattened map of the mouse cortex. The flatmap is generated by constructing a 3D-to-2D mapping such that the 2D Euclidean distance of every point on the flatmap to a pair of anchor points are the same as their 3D geodesic distance (shortest path along surface), resulting in the coordinate along one axis formed by the anchor points. This process is repeated for a second pair of anchor points to form the second axis. The white dotted line indicates the boundaries of what is visible in the top-down view in **b**. **(d)** The positions of all 1,082 injection centroids are plotted on the flat cortical surface. The 850 experiments are color coded as indicated by layer-specific projection class. The remaining 232 are shown in dark gray.

Across the 50 mouse lines (49 Cre and wild type), we generated a total of 1,082 experiments (**Supplemental Tables 1 and 2**). Most of these experiments (n=850) used 15 out of 50 lines (14 Cre and wild type). Figure 1a shows the 850 tracer experiments by line and cortical area. Mouse isocortex is parcellated into 43 areas in the Allen CCF, visualized here in two ways. A top-down view (Figure 1b), and a flattened view of the entire right hemisphere of cortex, to show all cortical areas in one 2D image (Figure 1c). Locations of the injections’ centroids (x,y,z voxel coordinate after registration to the Allen CCF) for all 1,082 experiments are plotted onto the cortical surface flat map in Figure 1d. Left side injections were plotted here on the right hemisphere to visualize completeness of areal coverage. All subsequent analyses refer to “ipsilateral” and “contralateral” relative to the side of injection. Of the 43 cortical areas, only 10 had 5 or fewer experiments. These areas (FRP, AIv, SSp-un, AIp, GU, VISC, AUDv, TEa, PERI, and ECT) were generally harder to target due to their size (SSp-un) or their location in very ventral or lateral regions (see Figure 1c). Abbreviations for isocortex regions are included in **Supplemental information**.

Data from all 1,000+ experiments in the isocortex are publicly available at the Allen Mouse Brain Connectivity Atlas data portal (http://connectivity.brain-map.org/). Individual experimental IDs and associated metadata are listed in **Supplemental Table 2.**

We visually inspected the brain-wide axonal projection patterns and classified all 1000+ experiments based on the principles described above for defining IT, PT, and CT neuron classes. Each experiment was manually assigned to one of five groups (**Supplemental Figure 3a-e**); (a) IT PT CT, when labeled axons were observed in all regions of interest (ipsilateral and contralateral cortex and striatum, and subcortical and thalamic projections), (b) IT, when labeled axons were restricted to ipsilateral and contralateral cortex and striatum, (c) PT, when labeled axons were ipsilateral and subcortically-projecting, (d) CT, when labeled axons projected almost exclusively to thalamus, and (e) local, when no (or few) long-distance (*i.e.,* outside of the source area) axons were seen. The consensus of results across all sources for each mouse line is shown in the “projection class row” of Figure 1a. More details can be found in **Supplemental Figure 3f**, which shows both the consensus projection neuron class for each Cre line and the class per source mapped within a Cre line. A subset of Cre lines are highly selective for IT, PT, or CT neurons, consistent with previous characterizations^22,29,30^. Most lines label neurons of the same projection class independent of the source area, but there are interesting exceptions. For example, L5 cells expressing Cre in the Chrna2-Cre_OE25 line are of the PT class in 14 of the 19 sources tested, but only locally projecting neurons are labeled in other sources (e.g. VISp) in this line. The manual assignment of experiments and Cre lines to projection neuron classes was also validated through unsupervised hierarchical clustering analysis of informatically-derived whole brain projection volumes in relevant major brain divisions (**Supplemental Figure 4**).

Together, the characterization of layer- and projection neuron class-selectivity for each Cre line enabled us to choose a core set of the best lines for comprehensively mapping connectivity from known classes of projection neurons in each cortical layer. These 13 lines, together with experiments in wild type mice (C57BL/6J) and the pan-layer Emx1-IRES-Cre line were used to identify all intracortical projections. These lines include L2/3 IT (Cux2-IRES-Cre and Sepw1-Cre_NP39), L4 IT (Nr5a1-Cre, Scnn1a-Tg3-Cre, and Rorb-IRES-Cre), L5 IT (Tlx3-Cre_PL56), L5 PT (A93-Tg1-Cre, Chrna2-Cre_OE25, Efr3a-Cre_NO108, Sim1-Cre_KJ18), L5-all classes (Rbp4-Cre_KL100), and L6 CT (Ntsr1-Cre_GN220, Syt6-Cre_KI148). One class for which we did not identify a suitable Cre line is L6 IT^16^.

### Intracortical connections are organized into modules

Cortical areas have distinct patterns (targets and weights) of corticocortical projections revealed through anterograde tracing in wild type mice^3,4^ (Figure 2). However, similarities between output patterns of some areas are also obvious when viewing the anatomical data spatially (Figure 2a) or the connection strengths in matrix form (Figure 2b). The matrix shows the output of a new data-driven model, which, for the isocortex, was based on 122 injections in wild type mice^31^. This model differs from our previously published model^4^ in that it is built through spatial interpolation at the voxel level (100 μm), rather than for each brain region, enabling the recovery of high spatial resolution for connectivity strengths between voxels. The voxel-based connection strengths were then unionized for every isocortical region annotated in the Allen CCF (n=43, Figure 1). Figure 2b shows the ipsilateral intracortical connectivity matrix.

**Figure 2.**
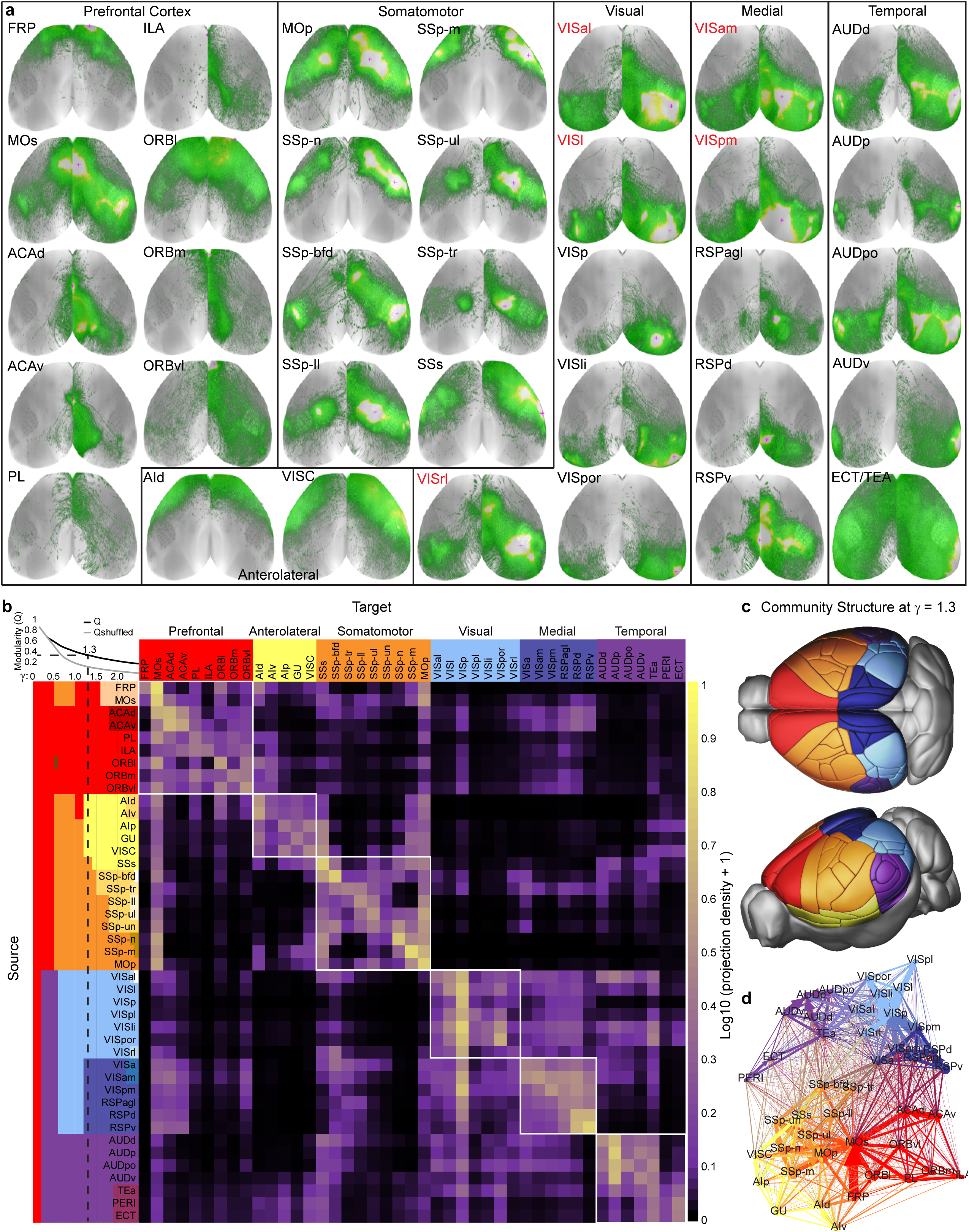
Modular organization of intracortical projection patterns based on the wild type connectivity matrix. **(a)** Top down cortical surface views showing the relative projection densities of labeled axons (normalized within each experiment, white is saturation) originating from 35 cortical source areas in C57BL/6J (black labels) or Emx1-IRES-Cre (red labels) mice. Red cross hairs indicate the location of the injection centroid. Some are not visible in the top down view. **(b)** Weighted connectivity matrix for 43 cortical areas. The data matrix was derived from the voxel-based model of Knox et al., 2018 and shows the connection strength as log10-transformed normalized projection density (the sum of predicted density per voxel in a target region normalized to that target’s volume). Rows are sources, columns are targets. Cortical areas are ordered first by module membership then by ontology order in the Allen CCF. Colors along the matrix axes indicate community structure with varying levels of resolution (*γ* = 0-2.5 on the y axis, *γ*= 1.3 only on the x-axis). The modularity metric (Q) is plotted for each level of *γ*, along with the Q value for a shuffled network containing the same weights. Community structure was determined independently for each value of *γ*, but colors were matched to show how communities split as the resolution parameter is increased. **(c)** Cortical regions color-coded by their community affiliation at *γ* = 1.3 show spatial relationships. **(d)** Diagram shows the ipsilateral cortical network in 2D using a force-directed layout algorithm. Nodes are color coded by module. Edge thickness shows relative projection density and edges between modules are colored as a blend of the module colors.

We analyzed the network structure of this ipsilateral matrix using the Louvain algorithm from the Brain Connectivity Toolbox^32^. This algorithm maximizes a modularity metric (Q) to identify groups of nodes (cortical areas) most densely connected to each other compared to a randomized network. Q quantifies the fraction of connections inside modules minus the fraction of connections expected inside the same modules if the network were connected randomly, *i.e.,* Q=0 has no more intramodule connections than expected by chance, while Q>0 indicates a network with some community structure.

To identify stable modules, we systematically varied the spatial resolution parameter, *γ*, from 0 to 2.5, and measured Q at each value of *γ*, compared to Q for a shuffled network. Increasing *γ* enables the detection of more modules, each containing fewer nodes^8,33^. The mouse cortex showed significant modularity (Q>Q for the shuffled network) for every value of *γ* above 0.3. Between 1 and 14 modules were identified across this range (Figure 2b, colors on left axis). For subsequent analyses, we chose to focus on the modules identified at *γ*=1.3 (Q=0.37). This value of *γ* corresponds to the midpoint between no modules at all, and the *γ* value of 2.5 where modules contain single regions. It is also the *γ* level where the difference between Q and Qshuffled was at its peak (0.2224±0.0021), although this difference was relatively stable between *γ*=1 and *γ*=1.8 (0.2187±0.0048 at *γ*=1, 0.2020±0.0009 at *γ*=1.8). The network is divided into six modules at this point, containing 5-8 regions each.

We named the six modules based on the cortical areas assigned to each; (1) **Prefrontal**: FRP, MOs, ACAd, ACAv, PL, ILA, ORBl, ORBm, ORBvl (2), **Anterolateral**: Aid, AIv, AIp, GU, VISC, (3) **Somatomotor:** SSs, SSp-bfd, SSp-tr, SSp-ll, SSp-ul, SSp-un, SSp-n, SSp-m, MOp, (4) **Visual**: VISal, VISrl, VISl, VISp, VISpl, VISli, VISpor, (5) **Medial**: RSPagl, RSPd, RSPv, VISa, VISam, VISpm, and (6) **Temporal**: AUDd, AUDp, AUDpo, AUDv, TEa, PERI, and ECT. Although we use these six modules for subsequent analyses, and to describe the organization of cortical areas based on connections, it should be emphasized that there are other, equally valid, levels of organization that could be chosen. For example, there is a four-module solution at *γ*=1.0 that results in prefrontal, somatomotor, visual, and temporal modules, in which the anterolateral areas are not split from somatomotor areas, and medial regions are still grouped with the visual areas. The spatial relationships between areas in these six modules are shown in the 3D renderings of brain areas in the Allen CCF in Figure 2c. There is a clear spatial component to the module assignment, in that nearby areas often belong to the same module. This is perhaps not surprising given that connectivity strengths drop as a function of distance^4,34^, but does not negate the likely importance of long-distance intermodule connections for integration and information flow. The network of connections within and between all modules and nodes was also visualized using a force-directed layout algorithm^35^ which highlights the overall high density, and variation in connection strengths across the cortex (Figure 2d).

### Interareal patterns of connectivity by output layer and class

Network analysis of the ipsilateral intracortical connectivity matrix revealed a modular organization based on the total output of a given cortical area. While this provides an important framework for understanding macroscale rules of cortical connectivity, the contributions of distinct cell classes within each area to the overall pattern are still unknown. To explore this, we first compiled groups of spatially-matched experiments. These experiments were pulled from the 850 listed in Figure 1a, using up to 15 mouse lines for coverage of layer/class within a given source. Each group was “anchored” by one of 90 Rbp4-Cre_KL100 tracer experiments (green triangles, Figure 1d). Rbp4-Cre_KL100 is a L5 selective line which labels all classes of projection neurons in L5. This line was chosen as the anchor because the largest number of sources were injected (33 of 43 cortical areas had at least 1 experiment) of all the Cre lines. Potential group members for each anchor were defined as experiments where the distance between the Rbp4 and other experiment injection centroids was <500 μm. An experiment was only used once, even if it was within 500 μm of two Rbp4 anchors. To be considered a complete group, at least one experiment from a Cre line representing L2/3 IT, L4 IT, L5 IT, L5 PT, L6 CT, and a wild type or Emx1-IRES-Cre dataset had to be present. Within a group, the median distance from the Rbp4-Cre_KL100 anchor was 296 μm. For some anchors, when a specific Cre line was otherwise missing, the injection centroid distance exceeded 500 μm (range 502-616 μm, 24/332 total experiments). All experiments within a group utilized the same tracer (*i.e.*, SypEGFP experiments were not grouped with EGFP experiments). In this way, we identified 43 anchor groups composed of unique sets of experiments (n=364 total), representing 25 of 43 potential source areas (**Supplemental Table 3**).

The locations of all anchor groups and individual experiments are shown mapped onto the flat cortical surface view in Figure 3a. Five examples are shown to illustrate layer selectivity, spatial matching of injections into different lines, and cortical projection patterns by line, including sources from the prefrontal cortex module (ACAd and MOs, Figure 3b,c), somatomotor module (SSp-m, Figure 3d), visual module (VISp, Figure 3e), and medial module (VISam, Figure 3f). 2D overlays of the injection sites confirmed the expected layer selectivity and relative size or proportion of cells labeled in these different experiments (Figure 3b’-f’, and see **Supplemental Figure 5** for individual panels). For example, prefrontal areas ACAd and MOs are both agranular structures (lacking L4), and the corresponding injections into the predominantly L4 IT Cre lines Scnn1a-Tg-3 or Nr5a1-Cre (colored in magenta) result in much sparser labeling than for areas with a large L4, such as primary somatosensory and visual areas (compare Figure 3b’-c’ with d’-e’). The distribution and density of cells labeled after viral infection of Cre-dependent reporter in each injection also closely matches expectations based on ISH characterization of tdTomato expression in Cre x Ai14 reporter lines for these regions^23^ (http://connectivity.brain-map.org/transgenic).

**Figure 3.**
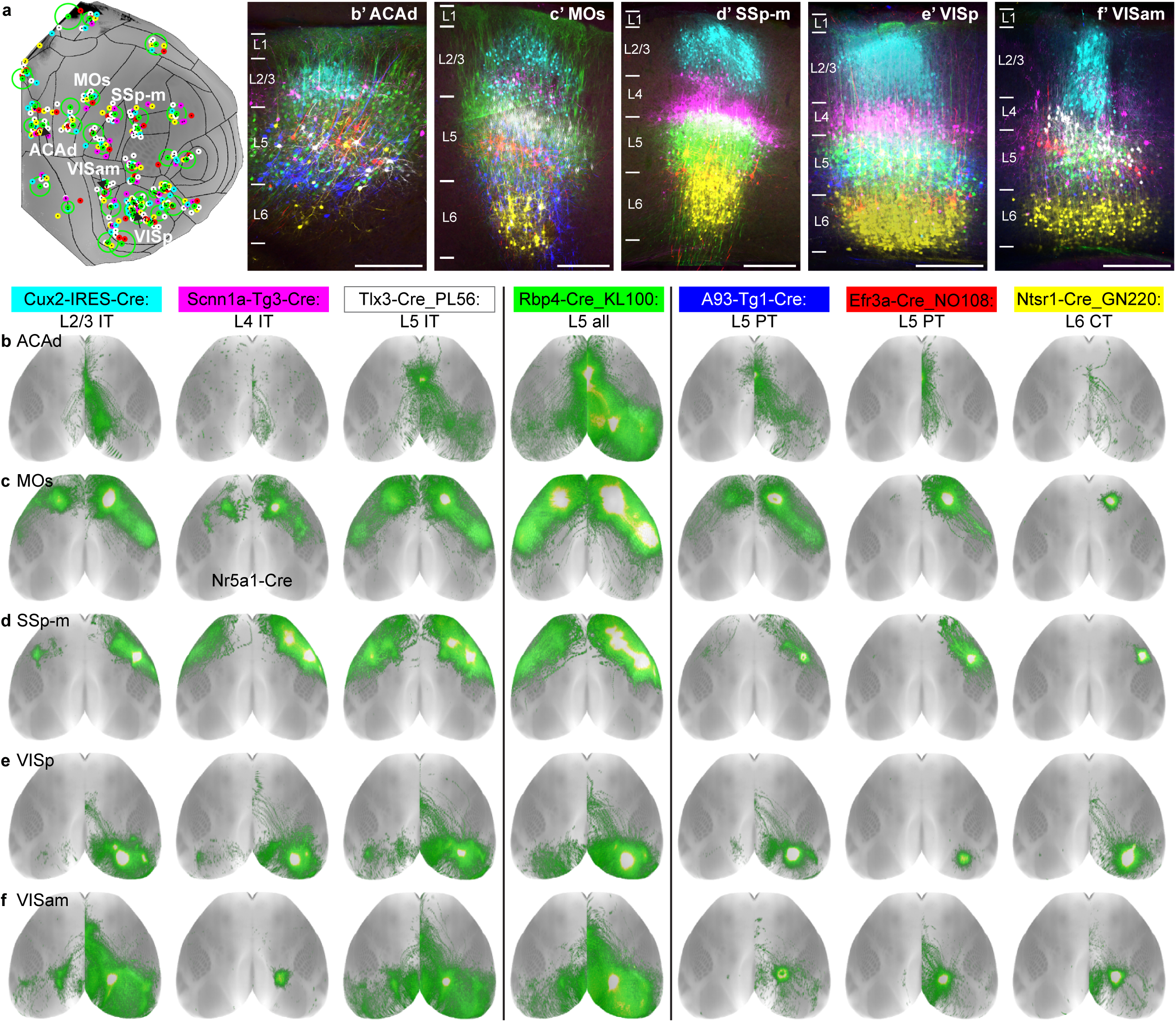
Comparison of layer- and class-selective intracortical projection patterns. **(a)** 43 groups of spatially-matched experiments to an Rbp4-Cre_KL100 anchor were collated based on having a “complete” membership roster; representing L2/3 IT, L4 IT, L5 IT, L5 PT, L6 CT and the L5 IT PT CT data from Rbp4-Cre_KL100. Each Rbp4-Cre experiment is shown as a green dot; all other experiments are color coded by layer and class as indicated. The green circle indicates the variance in distance to Rbp4 for each group. The five labeled groups are the examples shown in **b-f** (ACAd, MOs, SSp-m, VISp, and VISam). **(b’-f’)** 2-photon images acquired at the center of each injection site were manually overlaid by finding the best match between the top of L1 (pial surface) and bottom of L6 (white matter boundary) between each experiment, and then pseudocolored by Cre line to highlight the layer selectivity of Cre expression. Scale bar = 250 μm. **(b-f)** Top down views of the labeled axonal projections across the cortex originating from the infected neurons shown in **b’-f’**. Three Cre lines that label IT projection classes in L2/3 (Cux2-IRES-Cre), L4 (Scnn1a-Tg3-Cre, or Nr5a1-Cre as indicated for MOs) and L5 (Tlx3-Cre_PL56) are shown to the left of Rbp4-Cre_KL100. Three lines that predominantly label PT or CT projection neurons in L5 (A93-Tg1-Cre, Efr3a-Cre_NO108) and L6 (Ntsr1-Cre_GN220) are shown to the right. These lines also have intracortical projections, but target a smaller number of areas.

From any given source, cortical projections labeled in the L5 Rbp4-Cre_KL100 line (middle column in Figure 3b-f) appear to be more extensive than from any other line or layer. However, it is also visually obvious that the projections labeled in different Cre lines originating from the same location had very similar projection patterns overall (looking across rows). Indeed, it appears that the projections from every other spatially-matched Cre line are a subset of the L5 outputs. Of note, the predominantly subcortical projecting (see Figure 1b) L5 PT and L6 CT lines (3 columns on the right) also have varying amounts of intracortical projections, which are still a subset of those mapped from L5 IT and pan Cre lines.

To quantitatively explore how similar, or different, cortical projection patterns are across cell classes from the same location, we first manually curated the complete anchor group dataset (n=43 anchors, 364 experiments). This was accomplished by careful visual inspection of the corresponding high-resolution 2D images for all possible cortical targets to identify true positive and true negative connections from each experiment (43 ipsilateral and 43 contralateral targets, for a total of 31,304 connections manually checked, data provided in **Supplemental Table 3**). We also noted when a target contained only fibers of passage, and considered it as a true negative for subsequent binarization of the matrix. Using the output of our automated segmentation and registration algorithms we generated multiple weighted connectivity matrices, one for each Cre line, and applied the manually curated binary mask to remove all true negative weights (*i.e*., segmentation artifacts). As mentioned above, only 25 different cortical areas were represented in the 43 anchor groups. This was due to both denser spatial sampling within a larger structure (e.g. we targeted 6 retinotopic locations within primary visual cortex, 3 sub-regions in secondary motor and 2 in the ventral part of anterior cingulate), and the replication of experiments in several visual locations with the SypEGFP virus. To avoid biases related to differences between source areas, we selected only one anchor group per cortical region, if there was a significant, positive correlation between Rbp4-Cre_KL100 replicates (Spearman r >0.8). Following this selection, we present the results from 27 anchor groups, consisting of 25 unique areas and two locations in MOs and SSs. Eight of the lines representing each layer/class with the most experiments in these 27 groups are shown in Figure 4a (underlying data from these and the other 6 Cre lines provided in **Supplemental Table 3**). Of note, we merged the data from C57BL/6J and Emx1-IRES-Cre experiments into one matrix, as these both account for outputs of all projection neurons across layers in a cortical region. In support of this compilation, we also found that cortical projection patterns between pairs of spatially-matched Emx1-IRES-Cre and wild type experiments in 3 different regions were highly correlated, even given locations in opposite hemispheres (Spearman r= 0.88 for VISp, 0.90 for VISl, and 0.97 for VISam, **Supplemental Figure 6**).

**Figure 4.**
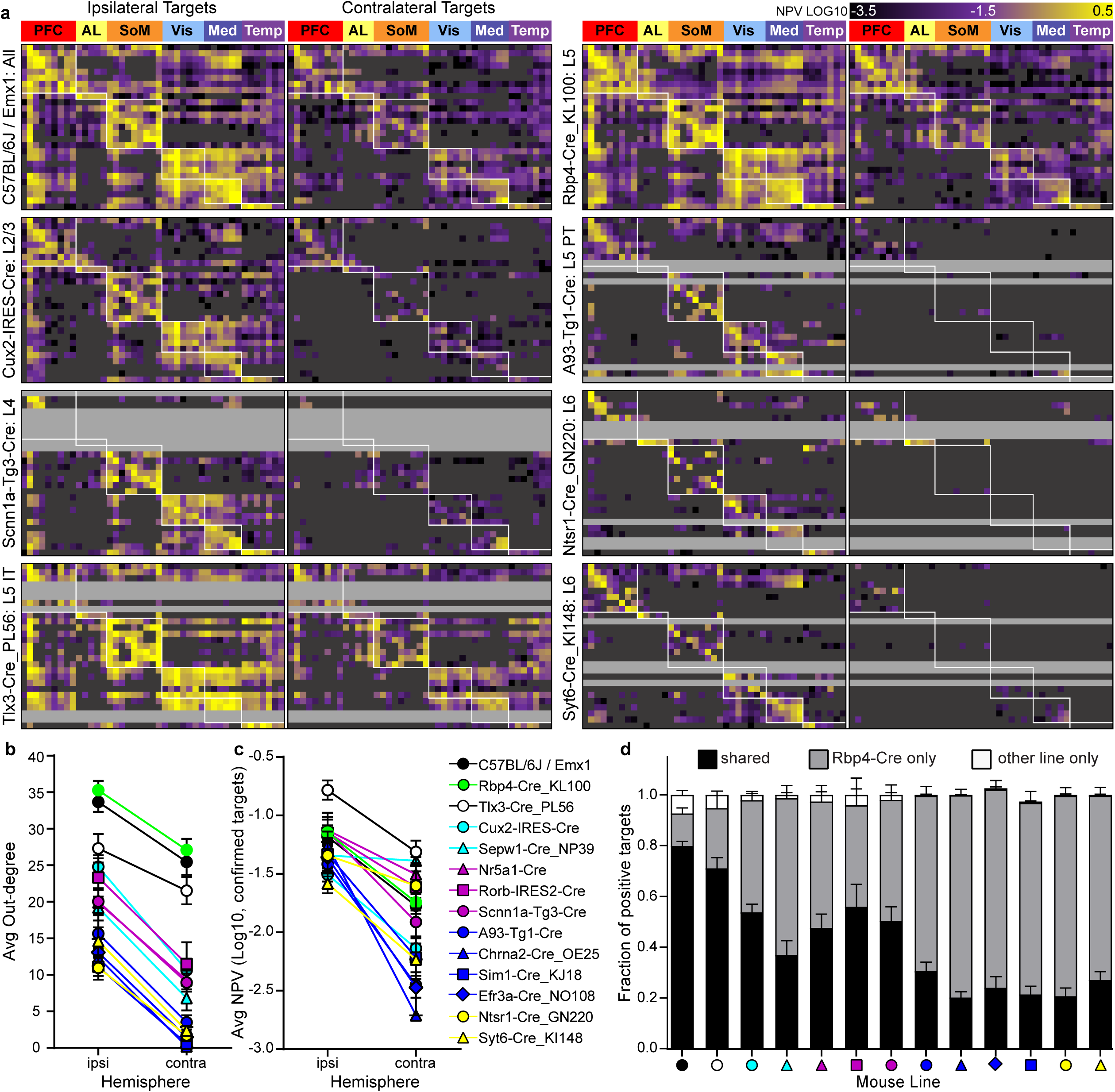
Cre line and layer-specific cortical outputs. **(a)** Eight directed, weighted, connectivity matrices (27 × 86) are shown for mouse lines representing projections labeled from different layers and cell classes. Each row of one matrix contains experimental data from one of 27 unique source areas. Columns show the 43 ipsilateral and 43 contralateral cortical target regions. Rows and columns follow the same module-based ordering in each matrix. Areas and connections belonging to the modules assigned using the ipsilateral voxel-based model data in Figure 2b are indicated by the white boxes. For every experiment, each of the 43 ipsilateral and 43 contralateral targets were inspected and assigned as containing either true positive or true negative axon terminal labeling. All true negatives (including passing fibers) were masked and colored dark grey. Rows for which an experiment was not completed are light grey. This was often because of low levels of Cre expression in those areas. The color map corresponds to log10-transformed normalized projection volumes in each target (range 10^−3.5^ to 10^0.5^, truncated at both ends). **(b)** The average out-degree across all sources represented in each matrix for each Cre line is plotted for the ipsilateral and contralateral cortex. **(c)** The average strength of all the connections (log10-transformed normalized projection volume) across source areas in each matrix are plotted for ipsilateral and contralateral hemisphere by Cre line. **(d)** Binary present or absent calls for the targets of each experiment were compared to the presence/absence calls from the matched Rbp4-Cre_KL100 anchor experiment. The average fraction of true positive targets shared by each line with its Rbp4 anchor experiment is plotted in the bar graph (black) as well as the average fraction of positive targets that are unique to Rbp4 (gray) or unique to the line indicated (white). Symbols or bars in **b-d** show mean +/- SEM.

Overall, these matrices reveal several similar and unique features of projection neuron class-specific connectivity between areas in terms of number, strength, and specificity of connections. First, we quantified the number of output connections (“out-degree”) for each experiment. Variation in out-degree by source area in each line is shown in **Supplemental Figure 7a**. To assess for differences across lines, we calculated the average out-degree in both hemispheres (Figure 4b). Overall, we find a significant effect of both Cre line and hemisphere (but not the interaction) on the number of connections (2-way ANOVA, p<0.0001). The mean numbers of C57BL/6J/Emx1-IRES-Cre connections are not significantly different from Rbp4-Cre_KL100 or Tlx3-Cre_PL56 in either hemisphere, but are significantly higher than all other Cre lines (Tukey’s multiple comparison test, p<0.0001), except for Rorb-IRES-Cre on the ipsilateral side. Similarly, Rbp4-Cre_KL100 also had significantly more connections on both ipsilateral and contralateral hemispheres compared to every other line, except for Tlx3-Cre_PL56 on the contralateral side. As also seen in the matrices, the L5 PT and L6 CT lines have the fewest number of connections to regions in both hemispheres, followed by the L2/3, L4, and L5 IT lines. Figure 4b also shows that for every line, there are fewer contralateral connections compared to ipsilateral connections.

Figure 4a and b show that L5 Rbp4-Cre_KL100 labeled cells project most widely, and are most like wild type experiments for any given source. Figure 3b-f also shows that the connections from each line appear to be a subset of the Rbp4-Cre_KL100 patterns, as opposed to a different set of target regions. So, next we determined how much overlap there was between the specific targets contacted by each experiment and the Rbp4 anchor within the spatially-matched groups (Figure 4d). C57BL/6J/Emx1-IRES-Cre and Rbp4-Cre_KL100 shared, on average, 80% of the targets from any given source. A roughly equal number of targets are unique to either Rbp4-Cre_KL100 or C57BL/6J/Emx1-IRES-Cre (12.7%, 7%) which may be due to differences in sensitivities of the viral tracers, or the homozygosity of the Emx1-IRES-Cre line. For every other Cre line, all target connections were a subset of the L5 Rbp4 targets (Figure 4c white bars, <5% of the targets are unique to any Cre line). Together, it appears that L5 cells project to almost all possible targets from any given source. Within L5, the L5 IT cells have the most overlap with Rbp4-Cre_KL100 while L5 PT cells have more limited projections within the Rbp4 set, and predominantly to the ipsilateral hemisphere (Figure 3b-f and 4a). L2/3 (and L4) IT cells project to a subset of the same targets of L5. Fewer projections to the contralateral hemisphere appear to account for most of the differences between L2/3 and L5 (Figure 4b), suggesting that most callosal projections arise from L5 in the mouse.

Next, we looked at the strength of connections made by the projection neurons labeled in each line (Figure 4c). After removal of the manually verified true negative connections, individual output strengths still spanned ∼5 orders of magnitude, like previously reported for both outputs and inputs in mouse and macaque brains^4,36,37^. Projection strengths were positively correlated (r=0.43 to 0.89) between all pairs of Cre lines from the same anchor group (**Supplemental Figure 8**). Overall, we find a significant effect of Cre line and hemisphere, and the interaction of these two factors, on the strength of connections (2-way ANOVA, p<0.0001 for Cre line and hemisphere, p=0.02 interaction effect). Across all lines, the average projection strengths are stronger within the ipsilateral compared to contralateral hemisphere (Figure 4c). This overall pattern was also observed across individual sources within each line (**Supplemental Figure 7b**). The largest disparities in strength across hemispheres is seen in the L5 PT (blue) and L6 CT lines (yellow). In other words, not only do these lines contact few targets contralaterally (<5, Figure 4b), but, when axon terminals are present, they are ∼1 order of magnitude weaker than the ipsilateral side connections.

### Axon terminal lamination patterns and their relationship to source layer and cell class

L5 neurons make connections to essentially all the targets of that source area, and all other layers contact a subset of these targets. Next, we looked at whether projection class-specific differences exist in the targets at the level of layers.

First, we visually inspected and described the relative densities of axon terminal labeling across layers for all targets in a subset of the Rbp4 anchor group experiments (79 of 364 experiments covering 14 source areas in all 15 lines; total=6,794 connections checked for layer patterns, true negatives then removed). Several frequently observed patterns emerged from this subset of representative experiments (Figure 5a-j). The four most common lamination patterns include: (**a**) columnar, with relatively equal densities across all layers (21%), (**b**) superficial and deep layers in equal densities (18%), (**c**) superficial layers only (25%), or (**d**) deep layers only (18%). The remaining patterns observed (**e-j**) together account for 19% of all true positive connections. Of note, almost all patterns involved dense terminals in L1, except for three rare patterns (**e, i, j**; 2-4%), which were distinctive in that L1 contained relatively few labeled axons.

**Figure 5:**
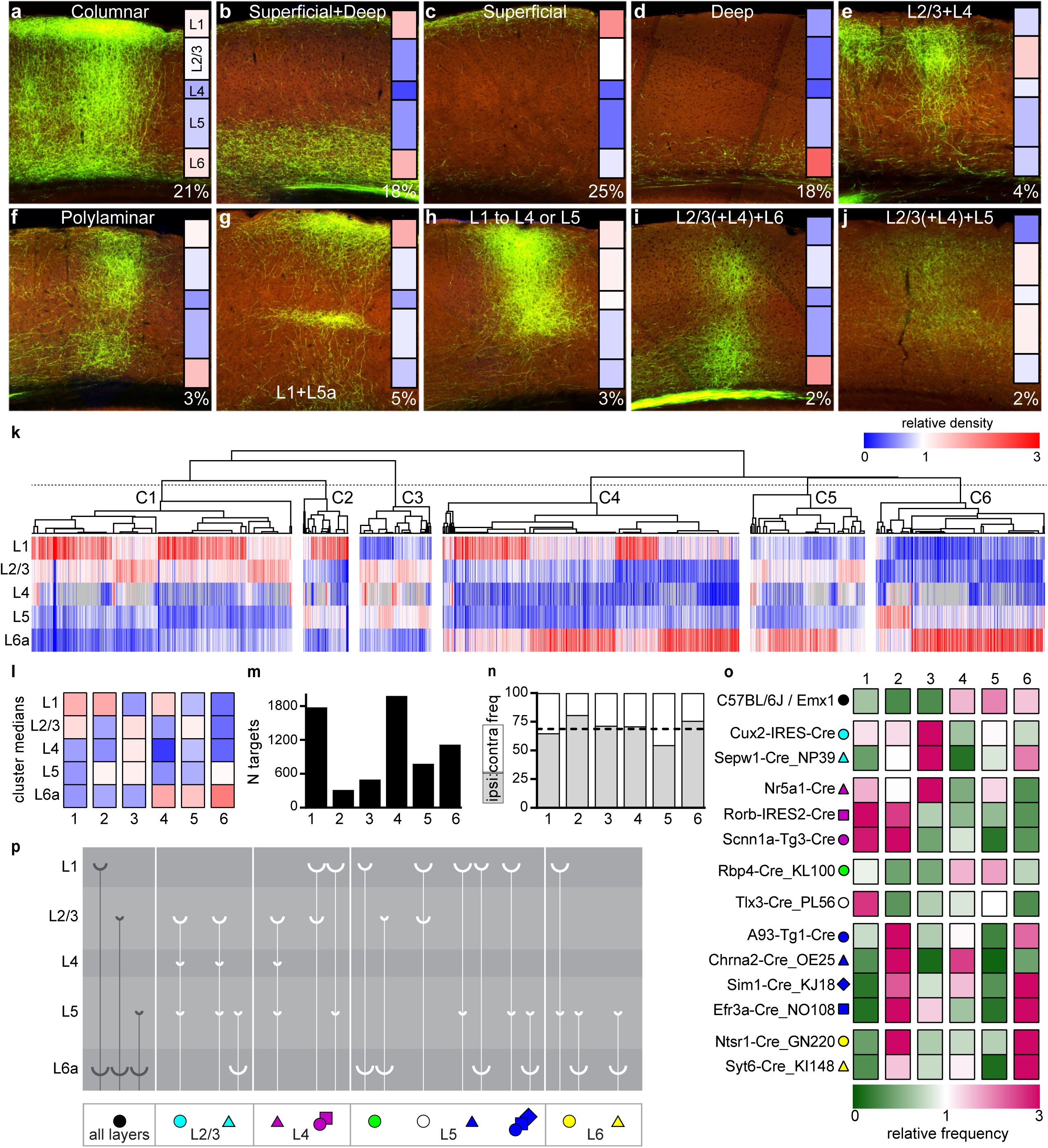
Diverse target lamination patterns in mouse cortex. **(a-j)** Relative densities of axon terminal labeling across layers for every cortical target were visually inspected for a subset of experiments, and then classified into one of ten categories based on overall observations. Four patterns occurred most frequently; **(a)** columnar, with relatively equal densities across all layers (21%), **(b)** superficial and deep layers in equal densities (18%), **(c)** superficial layers only (25%), or **(d)** deep layers only (18%). Additional patterns of note, although rare (<10%), included those in which L1 received relatively few axons **(e, i, j)**. Insets show the results of averaging informatically-derived quantification of relative layer density (the fraction of the total projection signal in each layer, scaled by the relative layer volumes) for all targets manually classified to that category. A relative density value of “1” (color = white) indicates that the fraction of axon labeling within a specific layer is equal to the relative size of that layer in that target, *i.e.,* it is neither more nor less dense than expected if axons were distributed evenly across layers, given differences in layer volumes. Values <1 indicate lower than expected density and >1 higher than expected density. Relative densities were color coded from 0 (blue) to 1 (white) to 3 (red). The color map key applies for panels a-l. **(k)** Unsupervised hierarchical clustering using spearman correlation and average linkages on the relative density values per layer. Each column is a unique combination of cre line, source area and target, after thresholding as described in the results. The dotted line indicates where the dendrogram was cut into 6 clusters. **(l)** Median relative density values by layer for each of 6 clusters. **(m)** Total number of targets in each cluster. **(n)** The frequency of ipsilateral and contralateral targets assigned to each cluster. The dotted line indicates the overall frequency of ipsilateral targets (68.77%). **(o)** The relative frequency of each Cre line appearing in one of the 6 clusters. The fraction of experiments in a cluster belonging to each Cre line was divided by the overall frequency of experiments from that Cre line in the complete dataset. A relative frequency value of “1” (color = white) indicates that Cre line appeared in that cluster with the same frequency as in the entire dataset. Values <1 (green) indicate lower than expected frequency, and >1 (pink) indicate higher than expected frequency of that Cre line in a cluster. **(p)** Schematic diagram showing the relationships between the layer and class of origin in the source area (Cre line symbols at the bottom) with the most frequent axon lamination patterns observed in the target area.

Following the qualitative assignment of axon terminals to a lamination pattern as shown, we checked whether informatically-obtained values of projection strength by layer, derived following registration to the Allen CCF, could quantitatively capture these patterns. The inset bars in Figure 5a-j show the average fraction of the total projection volume per layer, scaled by the relative size of the layer in each target. We used the actual ratio of each layer’s volume per target for scaling because this number varies across cortical areas (e.g. some areas have large L4, others very small). The resulting heat map in each panel visually corresponded well to the qualitative classifications of laminar patterns.

We next performed unsupervised hierarchical clustering, for the *complete* dataset of Figure 1a, to visualize laminar termination patterns from all source areas and Cre lines. In the heatmap shown in Figure 5k, each column is a unique combination of mouse line, source area and target. Relative density data were calculated as just described (*i.e.,* the fraction of the total projection in each layer, scaled by relative layer size). Data included for clustering had to pass three filters. (1) target connection strength (log10-transformed normalized projection volume) was greater than −1.5. This threshold was chosen based on the frequency distributions for informatically-derived normalized projection volumes of the set of manually-verified true positive and true negative connections (**Supplemental Figure 10**). At a log10 connection strength of −1.5, the number of true positives was first larger than true negatives. Less than 3% of true negative values remain, while over 50% of true positives are still present. (2) The percentage of infection volume in the primary source was > 50%, and (3) self-to-self projections were removed. Following these steps, if present, multiple experiments with the same source-line-target were averaged, resulting in a total of 6,469 unique source-line-target connections in Figure 5k.

We performed unsupervised hierarchical clustering on the relative density of projections in L1, L2/3, L4, L5, and L6a using Spearman rank correlations, and average linkages, to measure similarities. The first dendrogram branch point split the targets based on the density of projections to L6a (low on the left, high on the right). Then, within each of these two clusters, the next split was made by relative projection density in L1. The third split was determined by L2/3 relative projection density. At this point, 6 clusters were identified which resembled the manual categories, and we discuss each of these patterns further. The median values for each layer and the overall frequencies of these clusters are shown in Figure 5l,m. Clusters 1-3 had relatively weak projections to L6a compared to clusters 4-6. **(1)** Cluster 1 most resembled the superficial layer pattern (Figure 5c), with dense projections in L1 and/or L2/3 (n=1,777, 27%). **(2)** Cluster 2 resembled the L1+L5 pattern (Figure 5g; n=314, 4.9%). **(3)** Cluster 3 resembled two of the patterns avoiding L1 and L6 (Figure 5e,j, n=499, 7.7%), preferentially projecting into L2/3, L4 (if present), and L5. Unlike clusters 1-3, clusters 4-6 had high projection density to L6a. **(4)** Cluster 4 was the largest group (n=1,982, 31.0%) and most like the superficial and deep layer pattern (Figure 5b). Cluster 4 is also likely to contain the targets with lamination patterns visually described as “columnar” and “polylaminar” (Figure 5a,f), although even for those there does appear to be stronger projection density to L1 and L6 (see insets in a,f). **(5)** Cluster 5 targets were most densely innervated in L2/3 and L6a, like the pattern shown in Figure 5i (n=778, 12.0%). **(6)** Cluster 6 targets were most densely innervated in deep layers like in Figure 5d (L5 and L6a, n=1,119, 17.3%). All 6 of these broad classes of lamination patterns occurred in targets on the ipsi- and contra-lateral hemispheres at similar frequencies to the overall ratio of the number of ipsi- and contra-lateral targets (68.77% ipsilateral, Figure 5n).

Next, we wanted to determine the relationships (if any) between these laminar patterns and the Cre lines which label neurons of different layer-specific projection classes. For each mouse line, we calculated the relative frequency of that line in each cluster, divided by the overall relative frequency of each Cre line in the entire dataset. Figure 5o shows that the projections labeled in each mouse line have more than one type of target layer pattern (*i.e.,* very few of the boxes are 0, or colored dark green). However, for most lines, 1-3 layer patterns were identified that occur most frequently (pink-magenta). First, projections labeled following tracer injections into C57BL/6J and Emx1-IRES-Cre mice, which label the outputs of all layers and all classes together, are found with roughly equal frequencies in clusters 4,5, and 6 (all involving dense targeting to L6). The two L2/3 IT lines, Cux2-IRES-Cre and Sepw1-Cre_NP39 are most associated with cluster 3, as is one of the L4 IT lines, Nr5a1-Cre. In contrast, experiments from the other two L4 IT lines, Rorb-IRES-Cre and Scnn1a-Tg3-Cre, which had some selectively for L5 as well as L4 (**Supplemental Figure 2**) occur with higher than expected frequencies in clusters 1 and 2. The L5 pan-class line, Rbp4-Cre_KL100, is most associated with clusters 4 and 5, but the L5 IT line, Tlx3-Cre_PL56 is strongly associated with cluster 1. All four L5 PT lines were associated strongly with cluster 2, and three of these were also identified at higher than expected frequencies in cluster 6 (A93-Tg1-Cre, Sim1-Cre_KJ18, and Efr3a-Cre_NO108). The L5 PT line Chrna2-Cre_OE25, on the other hand, had relatively more projections of the cluster 4 type. Finally, L6 CT lines, Ntsr1-Cre_GN220 and Syt6-Cre_KI148, were like L5 PT lines in that they each had high relative frequencies of projections assigned to cluster 2 and 6 patterns.

The *most common* (but not all) laminar patterns from each Cre line are schematized in Figure 5p. In summary, L2/3 and L4 (Nr5a1) source neurons project predominantly to the middle layers in a target (L2/3, L4, and L5), avoiding L1. Other L4 source neurons project to L1 and either L2/3 or L5, avoiding L4 and L6. In L5, when both IT and PT classes are labeled, as in the Rbp4-Cre_KL100 line, projections target L6 and *either* L1 or L2/3. L5 IT source neurons predominantly target superficial layers (L1 and L2/3). L5 PT source neurons target either deep layers only (L5 and L6) or deep layers and L1, consistent with the L5 Rbp4-patterns representing both IT and PT patterns. L6 CT source neurons project to L1 and L5 or deep layers only.

### Anatomical rules for determining hierarchical position in mouse isocortex

Anatomical patterns of connections derived from anterograde and retrograde tracing data have been used to describe the hierarchical relationships between cortical areas for decades^12,26,38,39^. In the schemes based on the macaque monkey visual cortex, different lamination patterns correspond to feedforward, feedback, and lateral connections between pairs of areas. Briefly, feedforward connections were characterized by densest terminations in L4, feedback by the preferential avoidance of terminals in L4, usually with denser projections in both superficial and deep layers, and lateral connections characterized by having relatively equal density across all layers, including L4^12^. The fraction of retrogradely labeled cells in supragranular layers has also been used as a continuous variable index of hierarchical position in macaque cortex^39^.

Here, we observed some obvious similarities between the previously published laminar patterns derived from anterograde tracing data and our results, particularly for the feedback projection pattern. In our view, clusters 2 and 4 projection patterns (most like Figure 5g and Figure 5b, respectively) are most like the feedback rule described by Felleman and van Essen (1991). These patterns avoid L4, strongly targeting L1 and either L5 or L6 and can arise from neurons in all layers and classes (Figure 5o). However, we did not see a pattern emerge, either through unsupervised clustering or from the manual inspection, exactly like the feedforward projection rule of Felleman and van Essen (*i.e.,* preferentially targeting L4). The most similar projection pattern to a feedforward rule is seen in cluster 3 (most like Figure 5e,j), which involves projections into L2/3, L4, and L5, arising most often from L2/3 and one L4 IT class (Nr5a1 only, Figure 5o). The sparsity of projections into L1 with denser signal in mid layers is consistent with an index of feedforward connections recently described for mouse visual cortex^40^, and suggests that cluster 5 may also be a feedforward pattern in the mouse. Two patterns without an obvious match to previous literature were the superficial layer only projections in cluster 1 and the deep layer-only projections in cluster 6. Both patterns may be feedback because they do not involve mid-layers. Indeed, it was noted in Felleman and van Essen (1991) that the superficial only pattern was occasionally seen, and they grouped them with the feedback pattern because it did not involve L4. Also, of note, in the tracer experiments where all projection neuron classes were labeled (C57BL/6J/Emx1-IRES-Cre) the most common patterns were 4, 5, and 6. Based on the above descriptions, these would also support the identification of both feedforward (cluster 5) and feedback (cluster 4 and 6) types between areas when all neuron projection patterns between a given source-target are compiled.

To determine whether the tentative assignments of layer patterns to feedforward or feedback connections were consistent with past results and assumptions, we looked at specific pairs of connections where hierarchical relationships have previously been explored or intuited in rodents. Figure 6 shows two examples of reciprocally connected pairs of areas within unimodal sensory regions (visual and somatomosensory cortex) that are considered lower (primary) and higher (secondary) in a hierarchy. All projections out of VISp to higher visual areas are generally described as feedforward, whereas the reverse (to VISp from higher visual areas) are feedback^26,41–43^. We directly compared the axon projection patterns originating from neurons in different layers and classes in these examples using the spatially matched groups of experiments described in Figures 3 and 4. In the feedforward direction (VISp to VISal, Figure 6a), projections from VISp terminated with different layer patterns depending on the Cre-defined cells of origin. L2/3, L4, and L5 IT projections were densest in L2/3-L5 of VISal, and relatively sparse in L1 and L6. These connections were assigned to cluster 3. Rbp4-Cre_KL100 projections from VISp to VISal were densest in L2/3, L4 and L6, characteristic of cluster 5. The L5 PT and L6 CT cells projected to L1 and L5 (cluster 2). In the opposite direction (VISal to VISp, Figure 6a), patterns were often different from the corresponding reciprocal layer-specific projections. From VISal L2/3 IT cells, axons were distributed across all layers, with a sparser region in L5 (cluster 4). There was also a weak projection from L4 IT cells in VISal to VISp, with terminals in L1 and L5/6 (cluster 4). The projection originating from L5 IT cells ended predominantly in superficial layers (cluster 1), while the Rbp4-Cre_KL100 labeled axons from VISal to VISp were dense in L1 and deep layers (cluster 4). Again, projections from L5 PT and L6 CT cells were sparse, but present in both L1 and L6 (cluster 4). Overall, more of the projections in the feedforward direction involved middle layers, with sparser terminations in L1.

**Figure 6.**
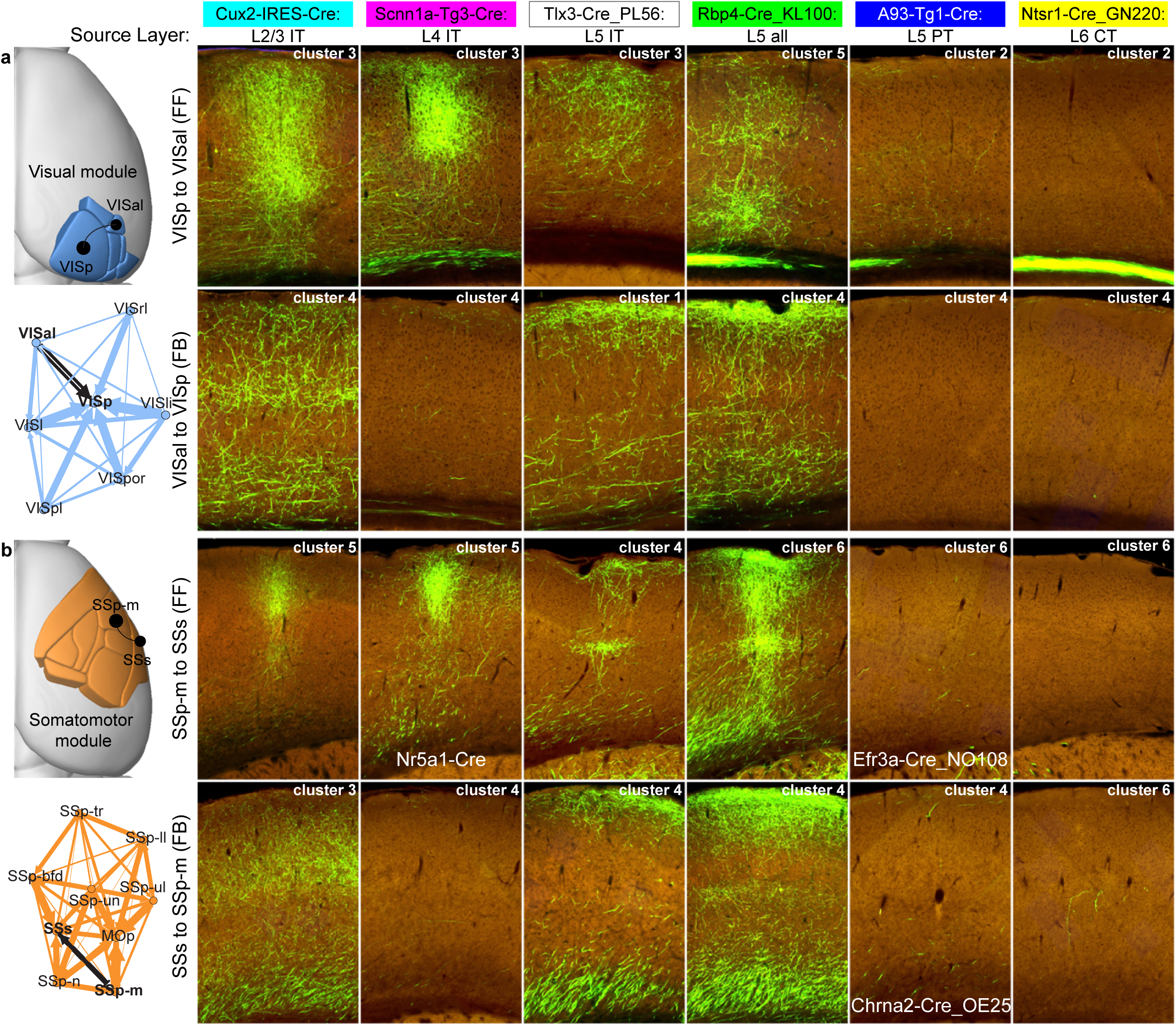
Intra-module projection patterns between reciprocally connected areas originating from different layers/classes. **(a)** In the visual module, VISp and VISal are reciprocally connected (black and white arrows). VISp is the de facto bottom of visual cortex hierarchies. The output to VISal from VISp is feedforward (FF). The reciprocal connection (VISal to VISp) is feedback (FB). **(b)** In the somatomotor module, the primary somatosensory cortex (SSp), like VISp, is the de facto bottom of the hierarchy. SSp-m sends feedforward projections to the secondary somatosensory region (SSs). SSs sends feedback projections to SSp-m. Modules from Figure 2b are shown spatially mapped on the cortex and as a force-directed network layout with the thickness of the lines corresponding to relative connection weights. 2P images in the approximate center of the axon termination fields for each target region show the laminar distribution of axons arising from labeled neurons in the different Cre lines, as indicated. Images were rotated so that the pial surface is always at the top of each panel. The cluster assignment for that line-source-target combination (columns **in** Figure 5k) is also indicated in each panel. One very striking difference between FF and FB connections was the strength and pattern of projections originating from L4 IT cells (second column). L4 IT cells in both modules strongly projected to the target in the FF direction, with patterns showing sparser axons in L1. In the FB direction, the L4 projection was weaker and ended in L1.

In the somatomotor module (Figure 6b), we focused on projections between a primary (SSp-m) and secondary (SSs) area as another example of a reciprocal feedforward (SSp-m to SSs) and feedback (SSs to SSp-m) connection. Like for the visual pair, projections from L2/3 and L4 IT cells preferentially innervated L2/3-L5, with relatively sparser terminals in L1 and L6. Both L5 IT and Rbp4-Cre_KL100 projections strongly innervate L1 and upper L5, unlike the VISp to VISal feedforward connection from L5 IT cells, which avoided L1. L5 PT and L6 CT cell projections were sparse, and to deep layers (cluster 6). In the reverse direction (SSs to SSp-m), the patterns looked remarkably like the layer-specific feedback projections from VISal to VISp. L2/3 IT cells terminated densely in mid layers (but appeared more diffuse across the entire column), but the other lines containing IT cells all had similar projections to superficial and deep layers. One very striking result from laying out the projection patterns originating from different layers is in the Scnn1a-Tg3-Cre: L4 IT column. For both examples (visual and somatomotor), there is a very strong connection originating from L4 cells in the lower area that preferentially terminates in mid-layers of the higher area. This is clearly not the case in the reverse direction (higher to lower area). Two additional examples of reciprocally connected areas within different modules (medial: RSPv to VISam and prefrontal: ORBl to MOs) are shown in **Supplemental Figure 11**. These generally follow the same patterns described above, including the obvious difference in strength and layer pattern in the L4 IT projection between reversed directions.

We next looked at the layer-specific projection patterns between reciprocally connected areas assigned to different network modules. The anterior cingulate cortex (ACA) exerts top-down control of sensory processing in VISp^44,45^. We thus assume that the intermodule connection from VISp to ACAd is feedforward, and ACAd to VISp is feedback (Figure 7a). In contrast to the intramodule feedforward connections in Figure 6 and **Supplemental Figure 11**, there is remarkable similarity in the target layer patterns arising from L2/3, L4, and all classes of L5 cells. These all preferentially innervate L1 in ACAd (cluster 1). In the feedback direction (ACAd to VISp), L2/3 cells also predominantly terminate in L1, but L5 cells project to both L1 and deep layers (L5 and L6, cluster 4), consistent with previous reports^44^. There may also be a sub-layer distinction in these L1 terminals. Axons from VISp to ACAd seem to be relatively deeper in L1 of ACAd, compared to the more superficial termination of ACAd axons in L1 of VISp. In Figure 7b, we present images showing laminar termination patterns arising from the different cell layer/classes between primary (MOp) and secondary (MOs) motor cortex, which we assigned here to the somatomotor (MOp) and prefrontal (MOs) modules based on their overall connectivity strengths with the other cortical areas. MOp is generally considered to be at a lower hierarchical level than MOs, although MOp is also the final output of the cortex driving voluntary control of behavior. All the IT cells from MOp to MOs appear to have more characteristics in common with intramodule feedforward connections (superficial layers, with sparse terminations in L1), including labeled axons from the predominantly L4 line. From MOs to MOp, there is more involvement of L1 in all patterns, and the L5 IT pattern includes deep and superficial layers like the Rbp4-Cre_KL100 experiment.

**Figure 7.**
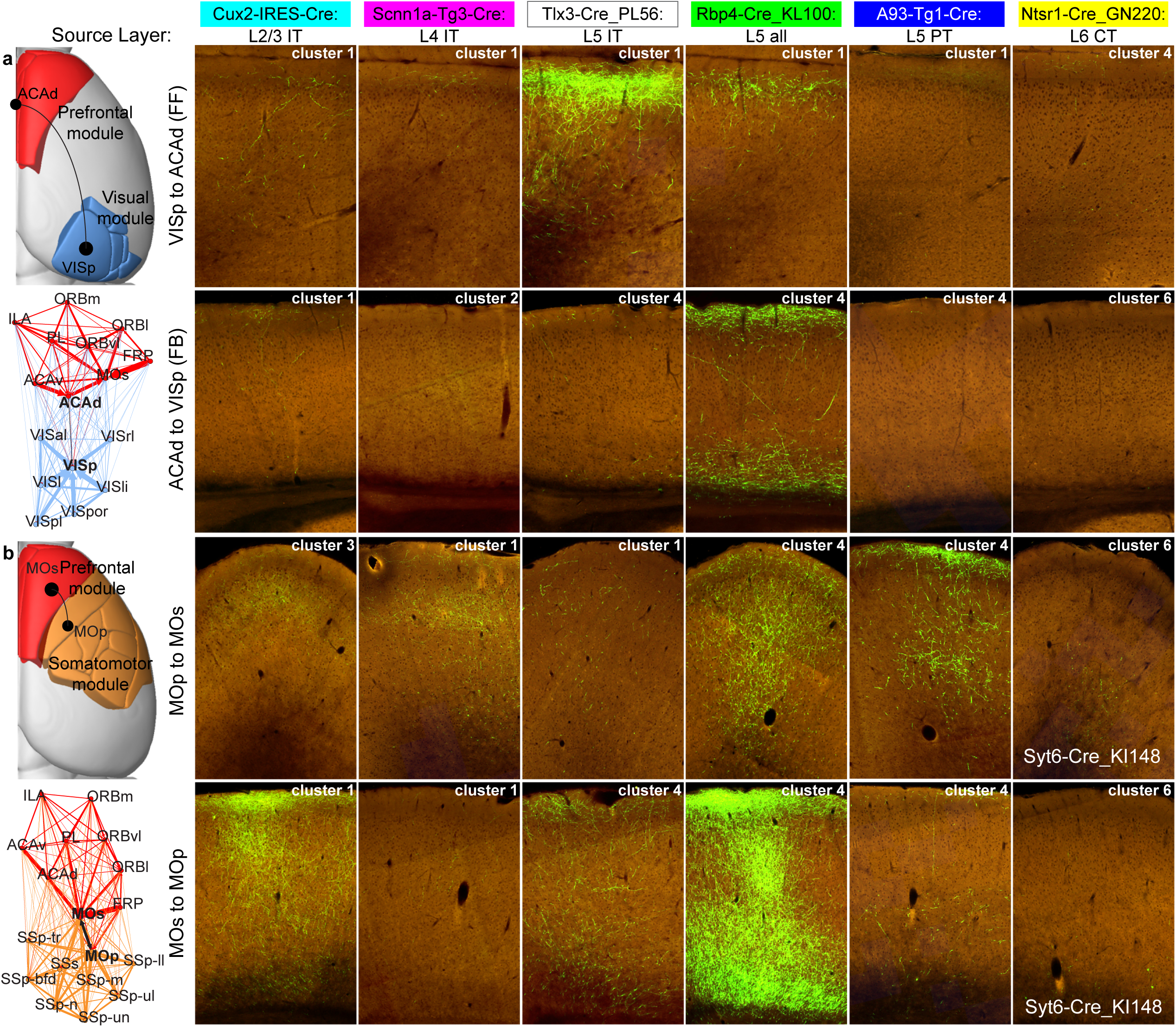
Inter-module projection patterns between reciprocally connected areas originating from different layers/classes. **(a)** Many reciprocal connections exist between areas in prefrontal and visual modules, e.g. VISp and ACAd. ACAd exerts top-down control of VISp activity, so we assume this connection is FB, and that the reverse is FF. **(b)** Reciprocal connections also exist between nodes of the prefrontal and somatomotor modules, e.g. MOs and MOp. Modules from Figure 2b are shown spatially mapped on the cortex and as a force-directed network layout with the thickness of the lines corresponding to relative connection weights. Like in Figure 6, 2P images in the approximate center of the axon termination fields for each target region show the laminar distribution of axons arising from labeled neurons in the different Cre lines, as indicated. Images were rotated so that the pial surface is always at the top of each panel. The cluster assignment for that line-source-target combination (columns **in** Figure 5k) is also indicated in each panel.

In summary, within a module, feedforward and feedback projections are consistent with the tentative assignments to cluster/layer pattern described above. Feedforward projections have more target patterns in clusters 3 and 5, feedback in clusters 2, 4 and 6. The relationship of cluster 1 to feedforward or feedback is less clear in these intramodule examples, although results in **Supplemental Figure 11** would support this pattern as feedforward, even with the L1 involvement. The intermodule connections from ACAd to VISp further support characterization of the cluster 4 layer pattern (superficial and deep, avoiding L4) as feedback, and, somewhat of a surprise, the cluster 1 pattern (superficial layers) as feedforward. Also, in all cases, projections from L5 PT and L6 CT neurons, when present, appear to be of the feedback type regardless of the overall top/down direction.

Taken together, these data suggest that the assignment of “feedforward” and “feedback” to specific connections between any pair of areas should account for all the contributions from each source layer, and the overall mix of target lamination patterns for a given connection (see also **Supplemental Figure 12** showing this concept, with more detail below).

Based on the anatomical analyses as described, we grouped the observed target layer patterns into either feedforward (clusters 1,3,5) or feedback (clusters 2,4,6) (Figure 8a). Quantification of the relative frequency of these patterns for intra- and inter-module connections showed that clusters 1 and 4 occurred more often than expected in intermodule connections; clusters 3, 5, and 6 were identified more often in intramodule connections (Figure 8b), consistent with data in Figure 6 and Figure 7. All source modules had relatively high frequencies of the feedforward pattern described by cluster 3 for their intramodule connections (Figure 8c). Between modules (Figure 8d), visual, temporal, and somatomotor were strongly associated with the superficial lamination pattern of cluster 1, which we classified as feedforward. Prefrontal connections were relatively more frequently assigned to cluster 4 (feedback), and all other modules occurred more often than expected in cluster 5 (feedforward).

**Figure 8.**
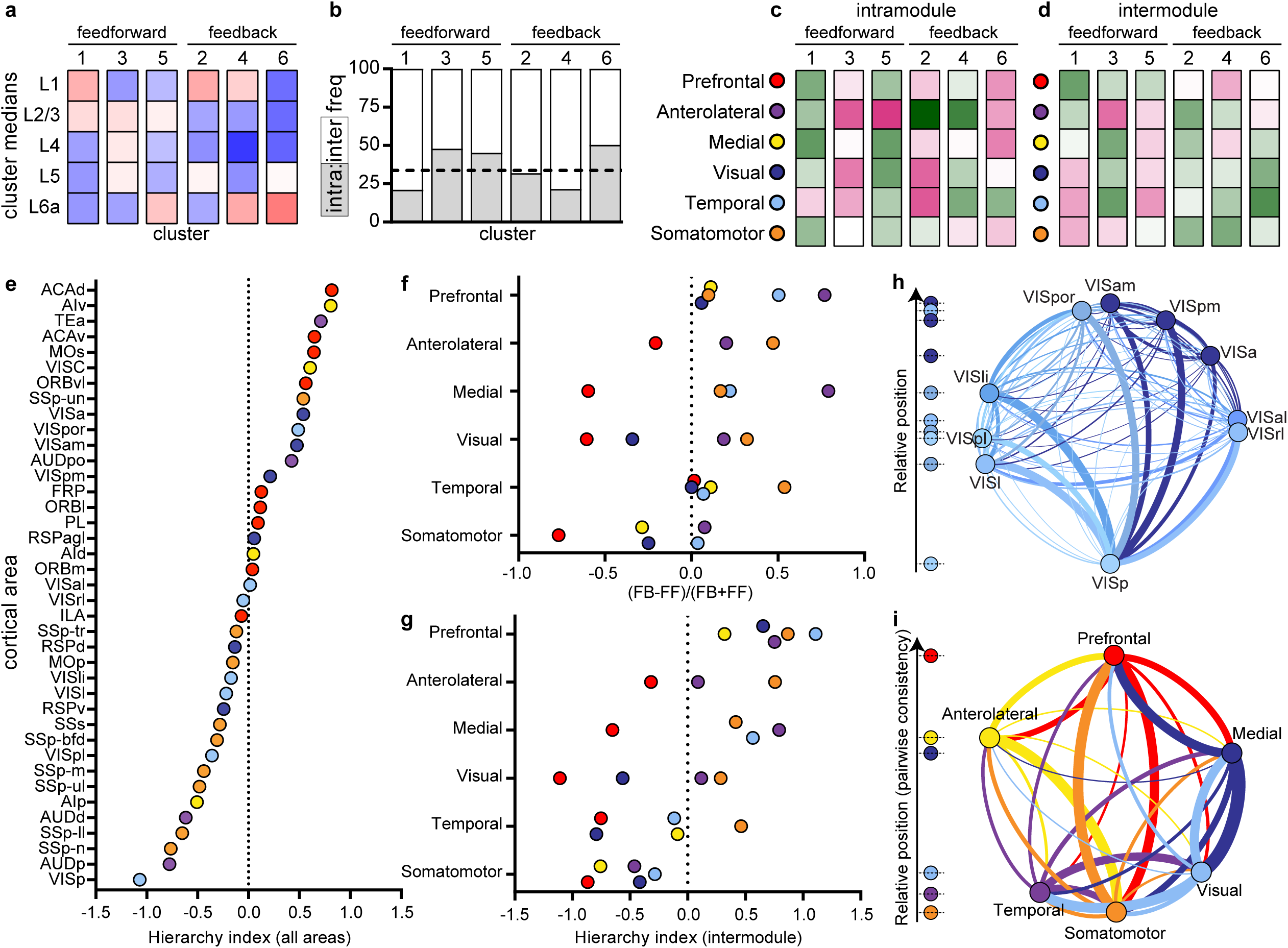
Organization of network modules into a hierarchy based on anatomical patterns of connections. **(a)** The six layer patterns identified through clustering in Figure 5k were classified as either feedforward or feedback. Clusters 1,3, and 5 were considered as characteristic of different feedforward projections. Clusters 2,4, and 6 were considered characteristic of different feedback projections. **(b)** The relative frequency of inter-module and intra-module connections is plotted for each cluster. The dotted line indicates the overall frequency of intramodule connections (33.74%). The laminar patterns of clusters 1 and 4 were relatively more frequent in intermodule connections, while clusters 3, 5, and 6 were associated more with intramodule connections. The relative frequencies within clusters are shown for each of the six modules as sources, for intramodule **(c)** and intermodule connections **(d)**. **(e)** 37 cortical areas are rank ordered by their hierarchical index scores and color coded by module assignment. **(f)** The relative differences in feedback to feedforward patterns between modules. Source module is indicated on the y-axis, and the relative differences in patterns between that module and every other target module is represented by the colored circles. Modules which had <10 connections were removed from analyses. Positive values indicate more feedback (FB) than feedforward (FF) connections from the source (y-axis) to target (plotted circles) module. Negative values indicate more FF than FB connection types. **(g)** The difference between the values plotted in (f) for each pair of modules as source and target, used to determine pairwise hierarchical positions. Positive values indicate the overall direction is feedback, given the reciprocal connection, from the source module on the y-axis to the target module (circles). Negative values indicate a feedforward connection from source to target modules. All intermodule connections from the prefrontal cortex were feedback. All from the somatomotor module were feedforward. Thus, these two modules formed the top and bottom of an intermodule hierarchy. **(h)** Network diagram showing interconnections of all 10 visual areas (visual module = blue, medial module = purple). Edge width indicates relative connection strength (from Figure 2b). The direction of the curved line shows outputs (clockwise) and inputs (counter-clockwise) from each node. Nodes are positioned along a circle perimeter based on hierarchical score, with VISp at the bottom and VISam at the top. All areas have strong feedback to VISp. **(i)** The intermodule network diagram shows each module as a node, with edge thicknesses based on the sum of connection weights from Figure 2b. Based on the pair-wise calls from data in **(g)**, we propose a hierarchical order of network modules that is consistent across levels. At the bottom is somatomotor, temporal, and visual, then medial, anterolateral, and, at the top, prefrontal.

### Unsupervised hierarchy of all cortical areas from layer termination patterns

We next determined whether it was possible to use the layer termination patterns from the layer-specific projection neuron classes defined by the Cre lines to define a direction of information processing throughout ipsilateral cortex. We defined hierarchical position for a cortical source area as the number of feedback connections originating from this area minus the number of feedforward connections, divided by their sum. Each connection was also normalized by a confidence level for the Cre line in providing information about the direction (see equations 1 and 2 in the methods). Similarly, the position in the hierarchy as a target is the normalized number of feedforward minus feedback connections terminating in the target. Each of these measures is bounded between −1 to 1. We used the sum of these measures as the hierarchical position (see equation 3 in the methods, possible range then becomes −2 to 2). Individual pair-wise measures for each connection, based on the clustering assignments to target lamination pattern types, are shown in **Supplemental Figure 12.** This matrix represents the concept of a “multigraph”, in that there are many possible edges between two nodes (e.g. each of the 13 Cre lines), and all information is used when searching for the optimal hierarchical positions. We searched over all possible mappings between the layer pattern type (the 6 clusters above) and feedforward and feedback assignments, and checked the self-consistency for every assignment. For the whole cortex assignment, the highest self-consistency (see equation 4 in methods) was obtained when clusters 1,3 and 5 were assigned to one type, and 2, 4 and 6 to the opposite. These are the *same* classifications derived from the above anatomical analyses, providing an example of how the human brain is an excellent unsupervised hierarchy discoverer.

Hierarchical positions of 37 (of the 43 parcellated) cortical areas are presented in Figure 8e (AUDv, GU, PERI, and ECT did not have data that passed thresholding; VISC and SSp-un were additionally removed for having only n=2 connections from 1 Cre line). By this measure, primary visual cortex (VISp) is at the bottom and the dorsal part of anterior cingulate cortex (ACAd) at the top. Areas were color coded by module assignment, which reveals a general pattern for prefrontal areas (red) to be higher in the hierarchy, and unimodal sensory regions (VISp, SSp, AUDp) to be closer to the bottom. For the entire cortex, the global hierarchy score (see equation 4 in methods) is 0.128 (range 0 to 1). Therefore, while we find a global hierarchy, there are still many connections for which a hierarchical organization may not provide a full explanation. We performed the same analyses on intramodule connections separately. Here, we observed a hierarchy score of 0.51 for the temporal, 0.33 for the visual module, 0.31 for anterolateral, 0.12 for medial, 0.09 for somatomotor, and 0.03 for prefrontal, given the same pattern assignments to feedforward (1,3,5) or feedback (2,4,6). The range in values suggests that different schemes, or different assignments of connections to feedforward and feedback, may better describe organization of connectivity within different modules. We also determined the relative order of all 10 visual areas, which span two modules here (visual and medial, colored blue and purple, respectively). Previous analyses based on tracer experiments in the mouse have mapped these areas into ventral and dorsal streams^46^. A representation of their hierarchical positions overlaid onto a weighted network diagram is shown in Figure 8h (dorsal stream areas on the right, ventral on the left of the circle). The edge weights in the network diagram are derived from the matrix of Figure 2b, and nodes are ordered from bottom to top (VISp, VISl, VISpl, VISrl, VISal, VISli, VISa, VISpm, VISpor, and VISam).

### Hierarchy of cortical network modules

We also determined the overall “feedback-ness” of each module’s ipsilateral outputs (as opposed to the individual nodes/areas) by measuring the number of feedback minus feedforward patterns, divided by their sum for every module as a source to all other modules (Figure 8f). Then, we calculated the difference between these feedback fractions for every pair of modules in both directions to obtain a single measure (the intermodule hierarchy index) predicting the forward/back relationship between each pair (Figure 8g). From this plot, it is obvious that the prefrontal module is above all other modules (e.g., every circle plotted for prefrontal as a source on the y-axis is positive on the x-axis). Conversely, the somatomotor module is below all other modules. We thus anchored the hierarchy at the top and bottom with prefrontal and somatomotor modules. Figure 8i shows a network diagram with each module collapsed into a single node. Edges are the sum of all the weights between modules from Figure 2b. The modules were positioned from top to bottom based on the data points in Figure 8g. Remarkably, the order is self-consistent at all levels given the available data. We did not have enough data or connections between anterolateral and medial, and anterolateral and visual modules to confirm these positions, however these are also the weakest of the intermodule connections (thinnest lines in Figure 8i). The combination of layer/class-specific projection patterns between network modules thus enabled the prediction of a consistent hierarchical framework for cortical information processing, with three modules containing primary sensory regions at the bottom (somatomotor, temporal, and visual), progressing to higher levels with modules containing more associational areas (medial, anterolateral) and ending in the prefrontal cortex.

## Discussion

Here we used a genetic tracing approach, building on our previously established viral tracing, whole brain imaging, and informatics pipeline, to map projections originating from unique cell populations in the same cortical area. Two key features of our high-throughput connectivity mapping pipeline are the automated segmentation of fluorescent signal, from which we calculate measures of long-distance projection strength between areas, and the registration of every experiment to our fully annotated 3D mouse brain reference atlas, the Allen CCF. Jointly, these methods enabled a comprehensive yet detailed view of mesoscale cortical wiring patterns, and the derivation of several general anatomical rules of long-range intracortical connections. Specifically, we show: (1) network analysis of intracortical connectivity patterns reveals a modular organization of cortical areas, (2) L5 neurons in any given source area make the most connections, and neurons in L2/3, L4, and L6 project to a subset of these L5 targets, (3) intracortical target lamination patterns are diverse, but at a coarse-grained level are related to layer of origin and are like previously described anatomical rules for defining feedforward/feedback connections, (4) projections originating from specific source layers/classes and target layer patterns together can define a single direction of information flow (*i.e.,* a hierarchy) between cortical areas and between network modules, with primary sensory areas and related modules at the bottom, and prefrontal areas at the top.

Our previously generated whole brain mesoscopic connectome provided a comprehensive, directed, and quantitative connectivity map between areas of the mouse cortex^4^. In the current study, we used these data and a novel voxel-based model^31^ to provide a macroscale organizational framework for viewing cortical areas as networks of connections. Through community detection analysis of the ipsilateral intracortical connectome, we identified six modules. The **first** module (“prefrontal”) consisted of cortical areas in predominantly frontal, agranular regions strikingly similar to those recently proposed as the mouse prefrontal cortex^47^. Very broadly, the function of the prefrontal cortex is cognition, and it is likely that the connections into and out of this module enable the necessary information flow for incoming sensory and behavioral state input that influence voluntary control of behavior. Indeed, there is relatively strong output from this module to primary motor cortex (MOp, Figure 2b), and strong input to secondary motor cortex (MOs) from many regions in other modules. The **second** module (“anterolateral”) included all three agranular insular subregions, plus gustatory and visceral cortex, consistent with a role for insular cortex in integration of taste and body homeostasis/energy needs^48,49^. At lower levels of spatial resolution (*γ*<1) these areas were part of a larger sensory module together with our **third** module (“somatomotor”). The somatomotor module contained all the primary somatosensory regions, secondary somatosensory cortex, and MOp. All SSp divisions project to SSs and MOp, and to MOs, which is in the prefrontal module. Although the MOp is not sub-parcellated in the Allen CCF, our projection mapping data show that projection domains are preserved in MOp when originating from the different SSp domains (*i.e.,* there are specific terminal fields in MOp for barrel field, trunk, lower limb, upper limb, nose, mouth), as previously reported^3^. The **fourth** module (“visual”) contained primary visual cortex and six out of nine higher order visual areas. The remaining three visual areas (VISa, VISam, and VISpm) were grouped with the three retrosplenial cortex subregions (RSPagl, RSPd, RSPv) into the **fifth** module (“medial”). At lower spatial resolution, these two modules (visual and medial) consistently merged into one, reflecting the high connectivity strengths between all visual areas. However, the medial module areas are more strongly connected to prefrontal areas (particularly ACAd, ACAv, and ORBvl). The **sixth** module (“temporal”) contained both auditory sensory regions (AUDp, AUDd, AUDv, AUDpo) and associational cortical areas, perirhinal (PERI), ectorhinal (ECT), and temporal association cortex (TEa). This combination is the least intuitive of all these modules, and may be a consequence of limitations in the underlying dataset. Specifically, in only one experiment did we successfully label projections from ECT cortex (shown in bottom right panel, Figure 2a), and so the voxel-based model had very little information to use for prediction of connection strengths^31^.

Within these modules and areas, we identified generalizable layer-specific intracortical projection patterns. For a given cortical source area, L2/3, L4, L5 PT, and L6 CT cells project to a subset of the target regions contacted by L5 neurons. Notably, all the excitatory neuron classes we surveyed had intracortical projections labeled outside of the infection area, including the L4 IT and subcortical PT and CT projection neurons. This somewhat unexpected result could be caused by Cre lines with less than perfect specificity. However, we also found that when long-distance projections are present from PT or CT cells, they have distinct termination patterns compared to the IT lines. These projection patterns were consistent with characteristics of feedback pathways, even in connections that were overall feedforward. A feedback collateral arising from deep layer subcortical-projection neurons may thus be another generalizable feature of PT and CT intracortical axons^50^.

The strength and presence of projections between areas from the predominantly L4 Cre lines was also unexpected. Canonical circuits, mostly derived from primate and cat, do not include inter-areal L4 excitatory neuron projections^51^. The three Cre lines used here had varying degrees of selectivity for L4 expression (**Supplemental Figure 2**), with some expression in L5 for both the Scnn1a-Tg3-Cre and Rorb-IRES-Cre lines. These two Cre lines also contain cells classified into both L4 and L5a types based on transcriptomics; while the Nr5a1-Cre line appears most specific to L4 cell types^15^. Thus, it is difficult to definitively conclude that these inter-areal projections originate from L4 rather than L5. Differences in the number of connections and the specificity of these inter-areal connections suggest that if the origin is in L5, they are at least a unique subclass of projection neurons. It is also worth noting that although the prefrontal and some other areas in the cortex are considered “agranular”, *i.e.,* lacking a distinct L4, Cre expression is often still detectable, though much sparser, in the so-called L4 lines used here.

Classic definitions for PT and CT cell classes exclude contralateral cortical projections^20^, roughly consistent with our observations. However, our data also showed that for some source areas, particularly in the prefrontal module, PT and CT lines had labeled axons that crossed the callosum, terminating in a small number of contralateral cortical targets. Overall, most callosal projections were made by neurons in L5, and, to a lesser extent, L2/3 and L4 Cre lines, consistent with expectations from previous results^52^. The number of ipsilateral connections made by L2/3 IT cells was similar to L5 IT cells; however, there were significantly fewer connections on the contralateral side. A recent analysis of the rat macroscale cortical connectome identified a set of general rules regarding ipsilateral (associational) and contralateral (commissural) connections that are mostly consistent with our observations; namely that all cortical areas have more associational than commissural targets^8^. The mesoscale connectome data we present here reveals that some of these differences can be better explained at the level of cell class.

Cortical areas are densely interconnected, and projections arise from all layers. Here, we first present an organizational scheme, the network, that groups cortical areas based on the strength of their connections. This kind of network view of cortical organization presents a structural view of all possible paths of information flow between areas and modules, but does not impose a direction or order on that flow. Another very influential organization scheme is the cortical hierarchy. The existence of a hierarchy implies classifying inter-areal connection types into at least two general classes: feedforward or feedback. From the macaque brain, studies have demonstrated that specific anatomical projection patterns between areas are characteristic of either a feedforward or feedback connection^12,13^. These primate rules were applied to build the visual cortex hierarchy that has inspired multiple computational models of cortical function^12^. Several models assign feedforward connections for information processing, and feedback connections as carrying a learning signal^14,53^. Another popular model for cortical computations is predictive coding^54^, in which feedforward connections represent an error signal, while feedback connections represent predictions, and local circuits integrate them^55^.

Partial hierarchies of the visual cortex exist for rodents, also based on anatomical projection patterns from anterograde tracing studies^26,40,56^. Differences between those patterns used in the primate were noted, and re-classified for rodent. Specifically, feedforward connections were characterized by having less dense axon terminations in L1 compared to L2/3, but axon terminals still spanned L2/3 to L5 evenly^40^. Feedback avoided L4 (like for the primate), terminating most densely in L1 and L6. We noted the same kinds of patterns in visual cortex feedforward and feedback connections. Whether these patterns can be extended to other sensory, motor, and associational modules (including those with “agranular” cortex^57^) was less clear from previous studies. Our analysis within and between modules suggests that there are several patterns associated with feedforward and feedback connections, but that every pattern can be classified into one or the other type. Within most modules, feedforward has the characteristics described above (i.e., densest in L2/3-L5), but between modules the feedforward connections were either dense in L2/3 and L6, or preferentially terminated in superficial layers (L1-L2/3). Feedback patterns matched previous descriptions in that there was preferential termination in L1 + deep layers (L5 or L6), or deep layers only. One of the most unusual patterns, not reported previously to the best of our knowledge, that clearly differentiated known feedforward from feedback connections was the strong presence of axons originating from L4 and ending in L2/3-L5. This occurred only in the feedforward direction of reciprocally connected pairs, but in the reverse direction there was both fewer axons from L4 and, when present, they were associated with more feedback patterns (avoiding L4). We also observed a striking relationship between feedforward/feedback patterns with the cell layer/classes, namely, supragranular (L2/3 and upper 4) neurons have predominantly feedforward projections, whereas infragranular (L5 and L6) neurons have both feedforward and feedback projections. However, as we have already noted, these types and their relationship to cell classes are also dependent on the specific connection.

Using these rules for feedforward and feedback across all cortical areas, we observe that the global organization of cortical connections is consistent with a hierarchy, in which a bottom-to-top direction can be defined. However, we want to emphasize that the hierarchical position does not explain all the connections, as the pathways between cortical areas are complex. The model used for Figure 8e shows an optimized hierarchy, but it is not the only possible solution, akin to the differences described for determining hierarchical order in primate visual cortex using the fraction of supragranular projection neurons (SLN), instead of discrete levels as in the Felleman and Van Essen diagram^12,39^.

Given the number of different connection types arising from a single area, we believe that new computational models, containing more than feedforward and feedback connections between nodes, are needed. This may be especially true when moving beyond models of sensory processing in the cortex. We would like to emphasize and encourage the adoption of a multigraph view of connectivity, in which two areas can be connected by multiple edges; each edge having an associated weight, type and subtype. We challenge the theoretical community to expand computational algorithms beyond those focused on classical graph structure. Additional data types that could predict directionality in cortical organization may also be added to these connections in the future. For example, the ratios of specific interneuron types, systematically mapped across all cortical areas, has recently been related to hierarchical position in the mouse^58^.

The expansion of the Allen Mouse Brain Connectivity Atlas to include mapping of projections from genetically-identified cell classes represents a big step toward a true *meso*scale connectome. Here, we present the addition of ∼ 1,000 new experiments to our online resource (http://connectivity.brain-map.org/), but focus only on the analysis of intracortical projection data. However, the complete brain-wide projection patterns are also already available for interested researchers to pursue a multitude of questions and analyses, and new results incorporating subcortical inputs and outputs may alter our view of a hierarchy in interesting and important ways. Finally, one of the limitations inherent in the population-based mapping approach used here is that we are still missing information at a more fine-grained level of cell types. Recent efforts and future work will undoubtedly further subdivide these broad classes of pyramidal neurons, at the level of areas as well as layers and projections, into specific cell types using morphology, physiology, and transcriptomics^15,17^. Here, we focused only on broad classes to derive general patterns of mesoscale cortical connectivity, which will be instructive and informative for future connectome data from more refined cell types. However, future large-scale efforts aimed at mapping the projections of specific cell types rather than classes, and even single cells^59,60^ will no doubt reveal additional principles of cell type-specific connectivity across the brain, moving us even closer to a full mesoscale connectome.

## Acknowledgements

We thank the Animal Care, Transgenic Colony Management and Lab Animal Services teams for mouse husbandry and tissue preparation. We thank all the members of the Neurosurgery and Behavior team for viral injections, including those not listed as authors: N. Berbesque, N. Bowles, S. Cross, M. Edwards, S. Lambert, W. Liu, K. Mace, N. Mastan, C. Nayan, B. Rogers, J. Swapp, C. White, and N. Wong. We also thank H. Gu for cloning of the synaptophysin-EGFP viral vector, E. Lee, F. Griffin, and T. Nguyen for intrinsic signal imaging, and J. Royall and P. Lesnar for schematic figure preparation. This work was supported by the Allen Institute for Brain Science and, in part, by National Institutes of Health grants R01AG047589 to J.A.H and U01MH105982 to H.Z. We thank the Allen Institute founder, Paul G. Allen, for his vision, encouragement, and support.

## Author Contributions

Conceptualization: H.Z., J.A.H., S.M. Supervision: H.Z., J.A.H., S.M., A.B., L.N., N.G., P.A.G., J.L., S.A.S, J.W.P., A.J., C.K. Project administration: S.M., S.W.O., W.W. Investigation, validation, methodology and formal analyses: J.A.H., S.M., K.E.H., J.D.W, J.K., P.B., S.C., L.C., A.C., N.G., N.G., C.G., P.A.G., A.M.H., A.H., R.H., L.K., J.L., J.L., M.T.M., M.N., L.N., B.O., S.A.S., Q.W., A.W, H.Z. Data curation: J.A.H., K.E.H., J.D.W., P.B., S.C., A.H., B.O., W.W. Visualization: J.A.H., K.E.H., J.D.W., L.N., D.F., S.M., M.N. The original draft was written by J.A.H., with input from K.E.H., J.D.W, S.M., Q.W., P.A.G., C.K., and H.Z. All co-authors reviewed the manuscript.

## Methods

### Mice

Experiments involving mice were approved by the Institutional Animal Care and Use Committees of the Allen Institute for Brain Science in accordance with NIH guidelines. Sources of mouse lines are listed in Supplemental Table 1. Characterization of the expression patterns of Cre driver lines used in this study have previously been described ^23^. Links to image series data are available through the Transgenic Characterization data portal (http://connectivity.brain-map.org/transgenic). Cre lines were derived on various backgrounds, but the majority were crossed to C57BL/6J mice and maintained as heterozygous lines upon arrival. Tracer injections were performed in male and female mice at an average age of P56 + 10 days. Mice were group-housed in a 12-hour light/dark cycle. Food and water were provided ad libitum.

### Tracers and injection methods

rAAV was used as an anterograde tracer. For most regions, stereotaxic coordinates were used to identify the appropriate location for a tracer injection^61^. For a subset of experiments in the left hemisphere, we first functionally mapped the visual cortex using intrinsic signal imaging (ISI) through the skull, described below. A pan-neuronal AAV expressing EGFP (rAAV2/1.hSynapsin.EGFP.WPRE.bGH, Penn Vector Core, AV-1-PV1696, Addgene ID 105539) was used for injections into wildtype C57BL/6J mice (stock no. 00064, The Jackson Laboratory). To label genetically-defined populations of neurons, we used either a Cre-dependent AAV vector that robustly expresses EGFP within the cytoplasm of Cre-expressing infected neurons (AAV2/1.pCAG.FLEX.EGFP.WPRE.bGH, Penn Vector Core, AV-1-ALL854, Addgene ID 51502). or, a Cre-dependent AAV virus expressing a synaptophysin-EGFP fusion protein to more specifically label presynaptic terminals (AAV2/1.pCAG.FLEX.sypEGFP.WPRE.bGH, Penn Vector Core).

Functional mapping of visual field space by intrinsic signal optical imaging (ISI) was used in some cases to guide injection placement. Additional details of this procedure can be found online (http://help.brain-map.org/display/mouseconnectivity/Documentation?preview=/2818171/10813533/Connectivity_Overview.pdf). Briefly, a custom 3D-printed headframe was attached to the skull, centered at 3.1 mm lateral and 1.3 mm anterior to lambda on the left hemisphere. A transcranial window was made by securing a 7-mm glass coverslip onto the skull in the center of the headframe well. Mice were recovered for at least seven days before ISI mapping. ISI was then used to measure the hemodynamic response to visual stimulation across the entire field of view of a lightly anesthetized, head-fixed, mouse. The visual stimulus consisted of sweeping a bar containing a flickering black-and-white checkerboard pattern across a grey background^62^. To generate a map, the bar was swept across the screen ten times in each of the four cardinal directions, moving at 9° per second. Processing of sign maps followed methods previously described^63^, with minor modifications. Phase maps were generated by calculating the phase angle of the pre-processed DFT at the stimulus frequency. The phase maps were used to translate the location of a visual stimulus displayed on the retina to a spatial location on the cortex. A sign map was produced from the phase maps by taking the sign of the angle between the altitude and azimuth map gradients. Averaged sign maps were produced from a minimum of three time series images, for a combined minimum average of 30 stimulus sweeps in each direction. Visual area segmentation and identification was obtained by converting the visual field map to a binary image using a manually-defined threshold and further processing the initial visual areas with split/merge routine^63^. Sign maps were curated and the experiment repeated if; (1) <6 visual areas were positively identified, (2) retinotopic metrics of V1 were out of bounds (azimuth coverage within 60-100 degrees and altitude coverage within 35-60 degrees) or, (3) auto-segmented maps needed to be annotated with more than 3 adjustments. Each animal had 3 attempts to get a passing map.

All mice received one unilateral injection into a single target region. For injections using stereotaxic coordinates from bregma as a registration point, an incision was made to expose the skull and bregma was visualized using a stereomicroscope. A hole overlying the targeted area was made by first thinning the skull using a fine drill burr, then using a microprobe and fine forceps to remove the bone, revealing the brain surface. For ISI-guided injections, the glass coverslip of the transcranial window was removed by drilling around the edges and a small burr hole drilled, first through the Metabond and then through the skull using surface vasculature fiducials obtained from the ISI session as a guide. An overlay of the sign map over the vasculature fiducials was used to identify the target injection site. rAAV was delivered by iontophoresis with current settings of 3 μA at 7 s ‘on’ and 7 s ‘off’ cycles for 5 min total, using glass pipettes (inner tip diameters of 10–20 μm). Mice were perfused transcardially and brains collected 3 weeks post-injection for Cre mice.

### Serial two-photon tomography

Imaging by serial two-photon (STP) tomography (TissueCyte 1000, TissueVision Inc. Somerville, MA) has been described^4,27^, and here we used the exact same procedures as our earlier published studies^4,25^.

### Image data processing

STP images were processed using the informatics data pipeline (IDP), which manages the processing and organization of the image and quantified data for analysis and display in the web application as previously described^4,28^. The two key algorithms are signal detection and image registration.

The signal detection algorithm was applied to each image to segment positive fluorescent signals from background. Image intensity was first rescaled by square root transform to remove second-order effects followed by histogram matching at the midpoint to a template profile. Median filtering and large kernel low pass filter was then applied to remove noise. Signal detection on the processed image was based on a combination of adaptive edge/line detection and morphological processing. Two variations of the algorithm were employed, depending on the virus used for that experiment; one was tuned for EGFP, and one for SypEGFP detection. High-threshold edge information was combined with spatial distance-conditioned low-threshold edge results to form candidate signal object sets. The candidate objects were then filtered based on their morphological attributes such as length and area using connected component labelling. For the SypEGFP data, filters were tuned to detect smaller objects (punctate terminal boutons vs long fibers). In addition, high intensity pixels near the detected objects were included into the signal pixel set. Detected objects near hyper-intense artifacts occurring in multiple channels were removed. We developed an additional filtering step using a supervised decision tree classifier to filter out surface segmentation artifacts (**Supplemental Figure 9**), based on morphological measurements, location context and the normalized intensities of all three channels.

The output is a full resolution mask that classifies each 0.35 μm × 0.35 μm pixel as either signal or background. An isotropic 3-D summary of each brain is constructed by dividing each image into 10 μm × 10 μm grid voxels. Total signal is computed for each voxel by summing the number of signal-positive pixels in that voxel. Each image stack is registered in a multi-step process using both global affine and local deformable registration to the 3-D Allen mouse brain reference atlas as previously described^28^. Segmentation and registration results are combined to quantify signal for each voxel in the reference space and for each structure in the reference atlas ontology by combining voxels from the same structure.

### Creation of the cortical top-down and flattened views of the CCF for data visualization

A standard z-projection of signal in a top-down view of the cortex mixes signal from multiple areas. Visualizations of fluorescence in Figures 1–3 instead project signal along a curved cortical coordinate system that more closely matches the columnar structure of the cortex. This coordinate system was created by first solving Laplace’s equation between pia and white matter surfaces, resulting in intermediate equi-potential surfaces. Streamlines were computed by finding orthogonal (steepest descent) paths through the equi-potential field. Cortical signal can then be projected along these streamlines for visualization.

A cortical flatmap was also constructed to enable visualization of anatomical and projection information while preserving spatial context for the entire cortex. The flatmap was created by computing the geodesic distance (the shortest path between two points on a curve surface) between every point on the cortical surface and two pairs of selected anchor points. Each pair of anchor points form one axis of the 2D embedding of the cortex into a flatmap. The 2D coordinate for each point on the cortical surface is obtained by finding the location such that the radial (circular) distance from the anchor points (in 2D) equals to the geodesic distance that was computed in 3D. This procedure produces smooth mapping of the cortical surface onto a 2D plane for visualization. This embedding does not preserve area and the frontal pole and medial-posterior region is highly distorted. As such, all numerical computation is done in 3D space. Similar techniques are used for texture mapping on geometric models in the field of computer graphics^64^.

### Data availability

Data (high resolution images, segmentation, registration to CCF, automated quantification) are available through the Allen Mouse Brain Connectivity Atlas portal (http://connectivity.brain-map.org/). In addition to visualization and search tools available at this site, users can also download data using the Allen Brain Atlas API (http://help.brain-map.org/display/mouseconnectivity/API) and the Allen Brain Atlas Software Development Kit (SDK: http://alleninstitute.github.io/AllenSDK/connectivity.html). Through the SDK, structure and voxel-level projection data is available for download. Examples of code for common data requests are provided as part of the Mouse Connectivity Jupyter notebook to help users get started with their own data analyses.

### Network modularity analysis

The matrix of connection weights between cortical areas (Figure 2b) was obtained from a novel model^31^. Briefly, this model allows us to predict the structural connectivity strengths between any given brain region in the mouse at the scale of voxels. This model combines the information from the ‘wild type’ viral tracing experiments performed as a part of the Allen Mouse Brain Connectivity Atlas. It uses the spatial information given by distances to injection sites to infer a connectivity strength from a given voxel to every other voxel in the Allen CCF.

We analyzed the network structure of this graph using the Louvain Community Detection algorithm from the Brain Connectivity Toolbox (https://sites.google.com/site/bctnet/)^32,65^. The Louvain algorithm uses a greedy algorithm to define groups of nodes (modules) that are more connected to each other than they are to other nodes outside their module. We determined the modularity at various levels of granularity by varying the resolution parameter, *γ*, from 0-2.5 in steps of 0.1. For each value of *γ*, the modularity was computed 1000x and each pair of regions received an affinity score between 0 and 1. The affinity score is the probability of two regions being assigned to the same module weighted by the modularity score (Q) for that iteration, thereby assigning higher weights to partitions with a higher modularity score. Each region was assigned to the module with which it had the highest affinity, with the caveat that all structures within a module had an affinity score >= 0.5 with all other members of the module. For each value of *γ*, we also generated a shuffled matrix containing the same weights but with the source and target regions randomized. The modularity for the cortical and hippocampal matrix (Q) and the shuffled matrix (Qshuffled) were evaluated at each value of *γ*

### Clustering Analyses and Statistics

Unsupervised hierarchical clustering was conducted with the online software, Morpheus, (https://software.broadinstitute.org/morpheus/) for algorithms and for visualization of the dendrogram and heat maps. Log-transforms were calculated on all values after adding a small value (0.5 minimum of the true positive array elements) to avoid Log (0). Proximity between clusters was computed using average linkages with spearman rank correlations as the distance metric. The clustering algorithm works agglomeratively: initially assigning each sample to its own cluster and iteratively merging the most proximal pair of clusters until finally all the clusters have been merged. To compare distances between granular and agranular samples (those that lack a L4), the computation of the distance metric was restricted to the set of shared layer projection fractions. In other words, we used the set of projection fractions in all layers when evaluating granular-granular distances, whereas when evaluating agranular-agranular or agranular-granular cortex, we used the set of all layers except L4. The software program GraphPad Prism was used for statistical tests and generation of all graphs, and the software program Gephi was used for visualization and layout of network diagrams.

### Unsupervised discovery of hierarchy position

Following the classification of the laminar patterns in clusters, we use an unsupervised method to simultaneously assign a direction to a cluster type and to construct a hierarchy.

First consider a mapping function

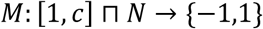

which maps a type of connection cluster to either feedforward (M=1) or feedback (M=-1) type. We search over the space of possible maps to see which map produces the most self-consistent hierarchy. Since some transgenic line have different numbers of connections in different clusters, some maps will lead to particular transgenic lines having very biased feedforward or feedback calls. Thus, we add a confidence measure, which decreases the importance of the information provided by a transgenic line to the global hierarchy if the calls from that transgenic line are biased.

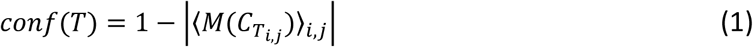

with a global confidence as an average over all the inter-areal connections above the threshold (10^-1.5)

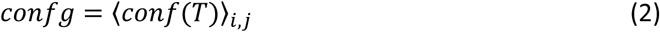

We define the hierarchical position of a source area based on the difference between the feedback and feedforward connections originating from this area, normalized by the number of connections, which is normalized by the confidence we have from different Cre lines providing information about the directionality of the connection. The hierarchical position as a target is the difference between the feedforward and feedback connections terminating in this area, normalized by the number of connections and confidence. The hierarchical position of an area is defined as the sum of these measures:

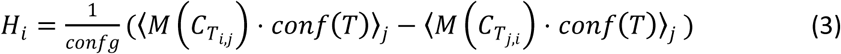

To test how self-consistent a hierarchy is we define the global hierarchy score:

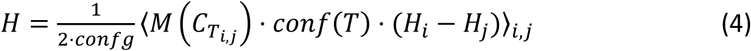

We performed an exhaustive search over all the maps M for the entire set of cortico-cortical connections, and the most self-consistent hierarchy is obtained when connections of type 1,3 and 5 are of one type and 2,4 and 6 are of the opposite type. Based on the position of the sensory areas, we conclude that type 1,3 and 5 are feedforward and 2,4 and 6 are feedback. It should be noted that a similar search inside of the visual module results in mapping connections 3 and 5 to feedforward and 1,2,4 and 6 to feedback.

## References

1. Sporns, O., Tononi, G. & Kötter, R. The Human Connectome: A Structural Description of the Human Brain. PLoS Comput. Biol. 1, e42 (2005).

2. Bohland, J. W. et al. A Proposal for a Coordinated Effort for the Determination of Brainwide Neuroanatomical Connectivity in Model Organisms at a Mesoscopic Scale. PLoS Comput. Biol. 5, e1000334 (2009).

3. Zingg, B. et al. Neural Networks of the Mouse Neocortex. Cell 156, 1096–1111 (2014).

4. Oh, S. W. et al. A mesoscale connectome of the mouse brain. Nature 508, 207–214 (2014).

5. Markov, N. T. et al. A weighted and directed interareal connectivity matrix for macaque cerebral cortex. Cereb. Cortex 24, 17–36 (2014).

6. Bota, M., Sporns, O. & Swanson, L. W. Architecture of the cerebral cortical association connectome underlying cognition. Proc. Natl. Acad. Sci. 112, E2093–E2101 (2015).

7. Scannell, J. W., Blakemore, C. & Young, M. P. Analysis of connectivity in the cat cerebral cortex. J. Neurosci. 15, 1463–83 (1995).

8. Swanson, L. W., Hahn, J. D. & Sporns, O. Organizing principles for the cerebral cortex network of commissural and association connections. Proc. Natl. Acad. Sci. 114, E9692–E9701 (2017).

9. Bullmore, E. & Sporns, O. Complex brain networks: graph theoretical analysis of structural and functional systems. Nat. Rev. Neurosci. 10, 186–198 (2009).

10. Sporns, O. Connectome Networks: From Cells to Systems. Micro-, Meso- and Macro-Connectomics of the Brain (Springer, 2016). doi:10.1007/978-3-319-27777-6_8

11. Wang, Q., Sporns, O. & Burkhalter, A. Network analysis of corticocortical connections reveals ventral and dorsal processing streams in mouse visual cortex. J. Neurosci. 32, 4386–99 (2012).

12. Felleman, D. J. & Van Essen, D. C. Distributed Hierarchical Processing in the Primate Cerebral Cortex. Cereb. Cortex 1, 1–47 (1991).

13. Rockland, K. S. & Pandya, D. N. Laminar origins and terminations of cortical connections of the occipital lobe in the rhesus monkey. Brain Res. 179, 3–20 (1979).

14. Riesenhuber, M. & Poggio, T. Hierarchical models of object recognition in cortex. Nat. Neurosci. 2, 1019–1025 (1999).

15. Tasic, B. et al. Adult mouse cortical cell taxonomy revealed by single cell transcriptomics. Nat. Neurosci. 19, 335–346 (2016).

16. Tasic, B. et al. Shared and distinct transcriptomic cell types across neocortical areas. bioRxiv 229542 (2017). doi:10.1101/229542

17. Zeng, H. & Sanes, J. R. Neuronal cell-type classification: challenges, opportunities and the path forward. Nat. Rev. Neurosci. 18, 530–546 (2017).

18. Sorensen, S. A. et al. Correlated gene expression and target specificity demonstrate excitatory projection neuron diversity. Cereb. Cortex 25, 433–449 (2015).

19. Shepherd, G. M. G. Corticostriatal connectivity and its role in disease. Nat. Rev. Neurosci. 14, 278–291 (2013).

20. Harris, K. D. & Shepherd, G. M. G. The neocortical circuit: themes and variations. Nat. Neurosci. 18, 170–181 (2015).

21. Gong, S. et al. Targeting Cre Recombinase to Specific Neuron Populations with Bacterial Artificial Chromosome Constructs. J. Neurosci. 27, 9817–9823 (2007).

22. Gerfen, C. R., Paletzki, R. & Heintz, N. GENSAT BAC Cre-Recombinase Driver Lines to Study the Functional Organization of Cerebral Cortical and Basal Ganglia Circuits. Neuron 80, 1368–1383 (2013).

23. Harris, J. A. et al. Anatomical characterization of Cre driver mice for neural circuit mapping and manipulation. Front. Neural Circuits 8, 1–16 (2014).

24. Daigle, T. L. et al. A suite of transgenic driver and reporter mouse lines with enhanced brain cell type targeting and functionality. bioRxiv 224881 (2017). doi:10.1101/224881

25. Martersteck, E. M. et al. Diverse Central Projection Patterns of Retinal Ganglion Cells. Cell Rep. 18, 2058–2072 (2017).

26. Coogan, T. A., Burkhalter, A. & Martin, K. Hierarchical organization of areas in rat visual cortex. J. Neurosci. 13, 3749–72 (1993).

27. Ragan, T. et al. Serial two-photon tomography for automated ex vivo mouse brain imaging. Nat. Methods 9, 255–8 (2012).

28. Kuan, L. et al. Neuroinformatics of the allen mouse brain connectivity atlas. Methods 73, 4–17 (2015).

29. Kim, E. J., Juavinett, A. L., Kyubwa, E. M., Jacobs, M. W. & Callaway, E. M. Three Types of Cortical Layer 5 Neurons That Differ in Brain-wide Connectivity and Function. Neuron 88, 1253–1267 (2015).

30. Olsen, S. R., Bortone, D. S., Adesnik, H. & Scanziani, M. Gain control by layer six in cortical circuits of vision. Nature 483, 47–52 (2012).

31. Knox, J. E. et al. High resolution data-driven model of the mouse connectome. bioRxiv 293019 (2018). doi:10.1101/293019

32. Rubinov, M. & Sporns, O. Complex network measures of brain connectivity: Uses and interpretations. Neuroimage 52, 1059–1069 (2010).

33. Sporns, O. & Betzel, R. F. Modular Brain Networks. Annu. Rev. Psychol. 67, 613–640 (2016).

34. Ria Ercsey-Ravasz, M. et al. A Predictive Network Model of Cerebral Cortical Connectivity Based on a Distance Rule. Neuron 80, 184–197 (2013).

35. Jacomy, M., Venturini, T., Heymann, S. & Bastian, M. ForceAtlas2, a Continuous Graph Layout Algorithm for Handy Network Visualization Designed for the Gephi Software. PLoS One 9, e98679 (2014).

36. Gămănuţ, R. et al. The Mouse Cortical Connectome, Characterized by an Ultra-Dense Cortical Graph, Maintains Specificity by Distinct Connectivity Profiles. Neuron 97, 698–715.e10 (2018).

37. Markov, N. T. et al. Weight Consistency Specifies Regularities of Macaque Cortical Networks. Cereb. Cortex 21, 1254–1272 (2011).

38. Maunsell, J. H. & van Essen, D. C. The connections of the middle temporal visual area (MT) and their relationship to a cortical hierarchy in the macaque monkey. J. Neurosci. 3, 2563–86 (1983).

39. Markov, N. T. et al. Anatomy of hierarchy: Feedforward and feedback pathways in macaque visual cortex. J. Comp. Neurol. 522, 225–259 (2014).

40. D’Souza, R. D., Meier, A. M., Bista, P., Wang, Q. & Burkhalter, A. Recruitment of inhibition and excitation across mouse visual cortex depends on the hierarchy of interconnecting areas. Elife 5, e19332 (2016).

41. D’Souza, R. D. & Burkhalter, A. A Laminar Organization for Selective Cortico-Cortical Communication. Front. Neuroanat. 11, 1–13 (2017).

42. Marshel, J. H., Garrett, M. E., Nauhaus, I. & Callaway, E. M. Functional Specialization of Seven Mouse Visual Cortical Areas. Neuron 72, 1040–1054 (2011).

43. Huh, C. Y. L., Peach, J. P., Bennett, C., Vega, R. M. & Hestrin, S. Feature-Specific Organization of Feedback Pathways in Mouse Visual Cortex. Curr. Biol. 28, 114–120.e5 (2018).

44. Zhang, S. et al. Selective attention. Long-range and local circuits for top-down modulation of visual cortex processing. Science 345, 660–5 (2014).

45. Leinweber, M., Ward, D. R., Sobczak, J. M., Attinger, A. & Keller, G. B. A Sensorimotor Circuit in Mouse Cortex for Visual Flow Predictions. Neuron 95, 1420–1432.e5 (2017).

46. Wang, Q., Sporns, O. & Burkhalter, A. Network Analysis of Corticocortical Connections Reveals Ventral and Dorsal Processing Streams in Mouse Visual Cortex. J. Neurosci. 32, 4386–4399 (2012).

47. Carlén, M. What constitutes the prefrontal cortex? Science 358, 478–482 (2017).

48. Avery, J. A. et al. Convergent gustatory and viscerosensory processing in the human dorsal mid-insula. Hum. Brain Mapp. 38, 2150–2164 (2017).

49. Hanamori, T., Kunitake, T., Kato, K. & Kannan, H. Responses of Neurons in the Insular Cortex to Gustatory, Visceral, and Nociceptive Stimuli in Rats. J. Neurophysiol. 79, 2535–2545 (1998).

50. Veinante, P. & Deschênes, M. Single-cell study of motor cortex projections to the barrel field in rats. J. Comp. Neurol. 464, 98–103 (2003).

51. Douglas, R. J. & Martin, K. A. C. NEURONAL CIRCUITS OF THE NEOCORTEX. Annu. Rev. Neurosci. 27, 419–451 (2004).

52. Fame, R. M., MacDonald, J. L. & Macklis, J. D. Development, specification, and diversity of callosal projection neurons. Trends Neurosci. 34, 41–50 (2011).

53. Yamins, D. L. K. & DiCarlo, J. J. Using goal-driven deep learning models to understand sensory cortex. Nat. Neurosci. 19, 356–365 (2016).

54. Rao, R. P. N. & Ballard, D. H. Predictive coding in the visual cortex: a functional interpretation of some extra-classical receptive-field effects. Nat. Neurosci. 2, 79–87 (1999).

55. Cain, N., Iyer, R., Koch, C. & Mihalas, S. The Computational Properties of a Simplified Cortical Column Model. PLOS Comput. Biol. 12, e1005045 (2016).

56. Coogan, T. A. & Burkhalter, A. Conserved patterns of cortico-cortical connections define areal hierarchy in rat visual cortex. Exp. brain Res. 80, 49–53 (1990).

57. Shipp, S. The importance of being agranular: a comparative account of visual and motor cortex. Philos. Trans. R. Soc. B Biol. Sci. 360, 797–814 (2005).

58. Kim, Y. et al. Brain-wide Maps Reveal Stereotyped Cell-Type-Based Cortical Architecture and Subcortical Sexual Dimorphism. Cell 171, 456–469.e22 (2017).

59. Han, Y. et al. The logic of single-cell projections from visual cortex. Nature (2018). doi:10.1038/nature26159

60. Economo, M. N. et al. A platform for brain-wide imaging and reconstruction of individual neurons. Elife 5, (2016).

61. Franklin, K. B. J. & Paxinos, G. Paxinos and Franklin’s The mouse brain in stereotaxic coordinates. (2012).

62. Kalatsky, V. A. & Stryker, M. P. New paradigm for optical imaging: temporally encoded maps of intrinsic signal. Neuron 38, 529–45 (2003).

63. Garrett, M. E., Nauhaus, I., Marshel, J. H. & Callaway, E. M. Topography and Areal Organization of Mouse Visual Cortex. J. Neurosci. 34, 12587–12600 (2014).

64. Oliveira, G. N., Torchelsen, R. P., Comba, J. L. D., Walter, M. & Bastos, R. Geotextures: A Multi-source Geodesic Distance Field Approach for Procedural Texturing of Complex Meshes. in 2010 23rd SIBGRAPI Conference on Graphics, Patterns and Images 126–133 (IEEE, 2010). doi:10.1109/SIBGRAPI.2010.25

65. Blondel, V. D., Guillaume, J.-L., Lambiotte, R. & Lefebvre, E. Fast unfolding of communities in large networks. J. Stat. Mech. Theory Exp. 2008, P10008 (2008).

